# Dynamics of oligomer populations formed during the aggregation of Alzheimer’s A*β*42 peptide

**DOI:** 10.1101/2020.01.08.897488

**Authors:** Thomas C. T. Michaels, Andela Šarić, Samo Curk, Katja Bernfur, Paolo Arosio, Georg Meisl, Alexander J. Dear, Samuel I. A. Cohen, Michele Vendruscolo, Christopher M. Dobson, Sara Linse, Tuomas P. J. Knowles

## Abstract

Oligomeric aggregates populated during the aggregation of the A*β*42 peptide have been identified as potent cytotoxins linked to Alzheimer’s disease, but the fundamental molecular pathways that control their dynamics have yet to be elucidated. By developing a general approach combining theory, experiment, and simulation, we reveal in molecular detail the mechanisms of A*β*42 oligomer dynamics during amyloid fibril formation. Even though all mature amyloid fibrils must originate as oligomers, we find that most A*β*42 oligomers dissociate to their monomeric precursors without forming new fibrils. Only a minority of oligomers converts into fibrillar species. Moreover, the heterogeneous ensemble of oligomeric species interconverts on timescales comparable to aggregation. Our results identify fundamentally new steps that could be targeted by therapeutic interventions designed to combat protein misfolding diseases.

---

Disorders of protein misfolding have recently emerged as some of the leading causes of death in the modern world.^1^ Over 50 such medical conditions have been identified, including Alzheimer’s disease, Parkinson’s disease, Huntington’s disease and amyotrophic lateral sclerosis, that are associated with the misfolding and subsequent aggregation of proteins into amyloid fibrils and plaques.^2–5^ Specifically, Alzheimer’s disease is linked to the self-assembly of the A*β*42 peptide and other length variants derived from the amyloid precursor protein, resulting in misfolded fibrillar aggregates that are observed as deposits in the brains of individuals suffering from this progressive disorder.^6,7^ The high molecular weight fibrils have been shown to be relatively inert, but lower molecular weight aggregates, denoted oligomers, have emerged as potent cytotoxins with the ability to trigger neuronal death in cell and animal models.^8–14^ New methods of kinetic analysis have transformed our understanding of the molecular events involved in the formation of the mature A*β*42 fibrils, revealing in particular that the proliferation of A*β*42 aggregates occurs at the surfaces of existing fibrils through an autocatalytic process known as secondary nucleation.^15–18^ Despite their fundamental importance, however, the detailed molecular mechanisms that drive the dynamics of cytotoxic oligomers during amyloid fibril formation remain unknown. In the present study, we have addressed this issue by providing direct measurements of the time evolution of oligomeric populations of A*β*42 formed during amyloid aggregation. These measurements, together with theory and simulations, define and quantify the fundamental molecular events of A*β*42 oligomer dynamics, providing new insights into the secondary nucleation step in A*β*42 aggregation.

## Results

We first obtained reproducible and quantitative measurements of A*β*42 oligomer concentrations formed during ongoing aggregation reactions starting from a supersaturated solution of ^3^H-labelled recombinant monomeric A*β*42.^19^ We then used centrifugation to remove fibrils, and utilised size-exclusion chromatography and liquid scintillation counting to identify the resulting oligomer fraction (Fig. 1 and Materials and Methods in Supplementary Sec. 1.6). This isotope-based approach is rapid, highly sensitive and does not rely on any form of chemical labelling, hence providing a method of studying oligomer populations quantitatively without perturbing their aggregation behaviour.^20,21^ ^3^H is used as a tracer replacing 0.1% of the protons bound to carbon, whereas all exchangeable and hydrogen-bonding positions contain ^1^H. As a validation of this approach we have obtained independent measurements of oligomer concentrations using mass spectrometry with a ^15^N isotope standard added after collecting the oligomer fractions (Fig. 1 and Supplementary Sec. 1.7). These measurements, shown in Supplementary Fig. 1, yield closely similar kinetics compared to the data measured using tritium labelling. Moreover, mass spectrometry is more sensitive than liquid scintillation counting, offering the additional possibility of quantifying the eluted fractions individually, hence allowing definition of the size distribution of the oligomer population (Supplementary Fig. 2).

**Figure 1:**
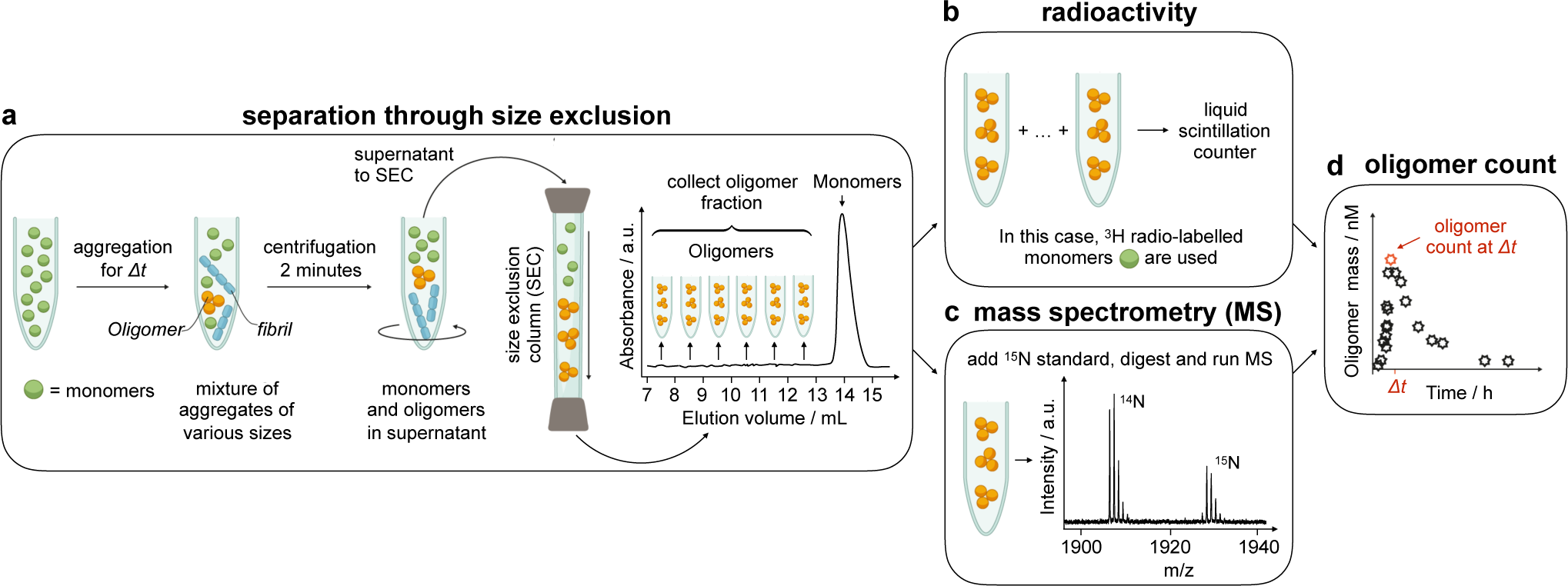
Experimental procedures for the quantitative measurement of A*β*42 oligomer populations during an ongoing amyloid fibril self-assembly reaction using tritium labelling or mass spectrometry. **(a)** We incubated varying concentrations of A*β*42 or A*β*40 monomers and collected aliquots at desired time points during the aggregation reaction. For each time point (∆*t*), we used centrifugation to remove fibrils. We then isolated the oligomeric fraction, encompassing species in the range of trimers to ca. 22-mers, through size-exclusion chromatography (SEC). We used a Superdex 75 column for which the void volume is ca. 7 mL and the monomer elutes at 14-16 mL. After separation through SEC, we used **(b)** liquid scintillation counting or **(c)** mass spectrometry (MS) to measure oligomer concentrations. In **(b)**, we used liquid scintillation counting to measure the absolute mass concentration of peptides eluting between 7 and 13 mL in the case of ^3^H-labelled A*β*42. In **(c)**, we used MS of natural abundance peptides, in which case each fraction (1 mL) was lyophilised, redissolved in 20 *µ*L H_2_O, supplemented by a defined amount of ^15^N-A*β*42 (10 pmol) and AspN enzyme, digested overnight, and analysed by MALDI-TOF MS. The peptide concentration in each fraction was determined as the ratio *r* of the integrated area of the ^14^N peak at 1906 m/z and the ^15^N peak at 1928 m/z as *c* = *r ×* 10 nM. The total oligomer concentration at each time point ∆*t* was calculated as the sum over fractions 7-12. The relative A*β* concentration in each fraction was then calculated by dividing *c* by the summed concentrations over fractions 7-12. **(d)** Observed concentration of oligomers versus aggregation time, ∆*t*. This procedure, which requires 16 minutes for oligomer isolation (Materials and Methods), provides a rapid and quantitative readout of the time evolution of oligomeric populations.

These oligomer population measurements were then combined with a rigorous kinetic analysis to develop a detailed mechanistic understanding of A*β*42 oligomer dynamics (Supplementary Secs. 3-6). In general terms, the formation of oligomers requires two or more monomers to come together, either through a primary (i.e. fibril-independent) or secondary (i.e. fibril-dependent) nucleation mechanism. The population of oligomers can then in turn decrease (i) through conversion of non-fibrillar to fibrillar oligomers, (ii) through the elongation of fibrillar oligomers, or (iii) through processes that do not lead to the oligomers being a source of new fibrils, such as dissociation into monomers. The population balance of each aggregate species can be parameterised mathematically through a master equation approach, similar to that developed for fibril formation,^15–18^ and we have exploited self-consistent approaches to develop integrated rate laws (Supplementary Secs. 4-5) to be compared directly with the experimental data discussed above (Supplementary Sec. 6).

Using this framework, we first addressed the fundamental question of whether or not the oligomers formed during the aggregation process are elongation competent, i.e. are able to sequester further monomers to grow in a manner similar to that established for mature amyloid fibrils.^16^ We measured the time course of the oligomer populations formed from a solution of monomeric A*β*42 at an initial concentration of 5 *µ*M and simulated a series of mechanistic scenarios to examine the quantitative level of agreement between the two. In the simplest scenario, all oligomers are short fibrillar aggregates with the same rate of elongation as mature fibrils (Fig. 2a). In this scenario, any decrease in the oligomer population can only arise through the direct growth of the oligomers into species which are larger than those in the oligomeric fraction captured in the experiment. The mathematical analysis of the master equation in this limit (Supplementary Sec. 4.3) shows that the oligomer dynamics are described by the same underlying microscopic parameters as those defining the kinetics of growth of the higher-molecular weight fibrils, which are available from the analysis of fibril mass kinetics recorded at varying monomer concentrations (Fig. 2a,i) (16). The kinetics of oligomer formation can therefore be computed directly, without any further free parameters, from the analysis of the bulk amyloid aggregation data. Comparison of the model predictions with the experimental data on oligomer concentrations reveals, however, that the latter exceed the theoretical predictions by 5 orders of magnitude (Fig. 2a,ii); the highest observed oligomer concentration over time is ca. 80 nM, while the model prediction is 280 fM, revealing that the model outlined in Fig. 2a is unable to describe, even approximatively, the observed behaviour. We thus conclude that A*β*42 oligomers measured in the experiments are structurally distinct from fibrillar aggregates in that they are unable to recruit monomers to grow in size as effectively as mature fibrils. Yet, all fibrils must originate from the growth of smaller oligomeric structures, implying that at least some oligomers must undergo a structural conversion to become faster elongating fibrillar structures. It is likely that such a structural reorganisation will occur during a conversion step to produce a fibrillar oligomer with very similar molecular packing to that observed in mature fibrils.^22,23^ The conversion process may occur in solution or in contact with fibril surfaces. ^24,25^

**Figure 2:**
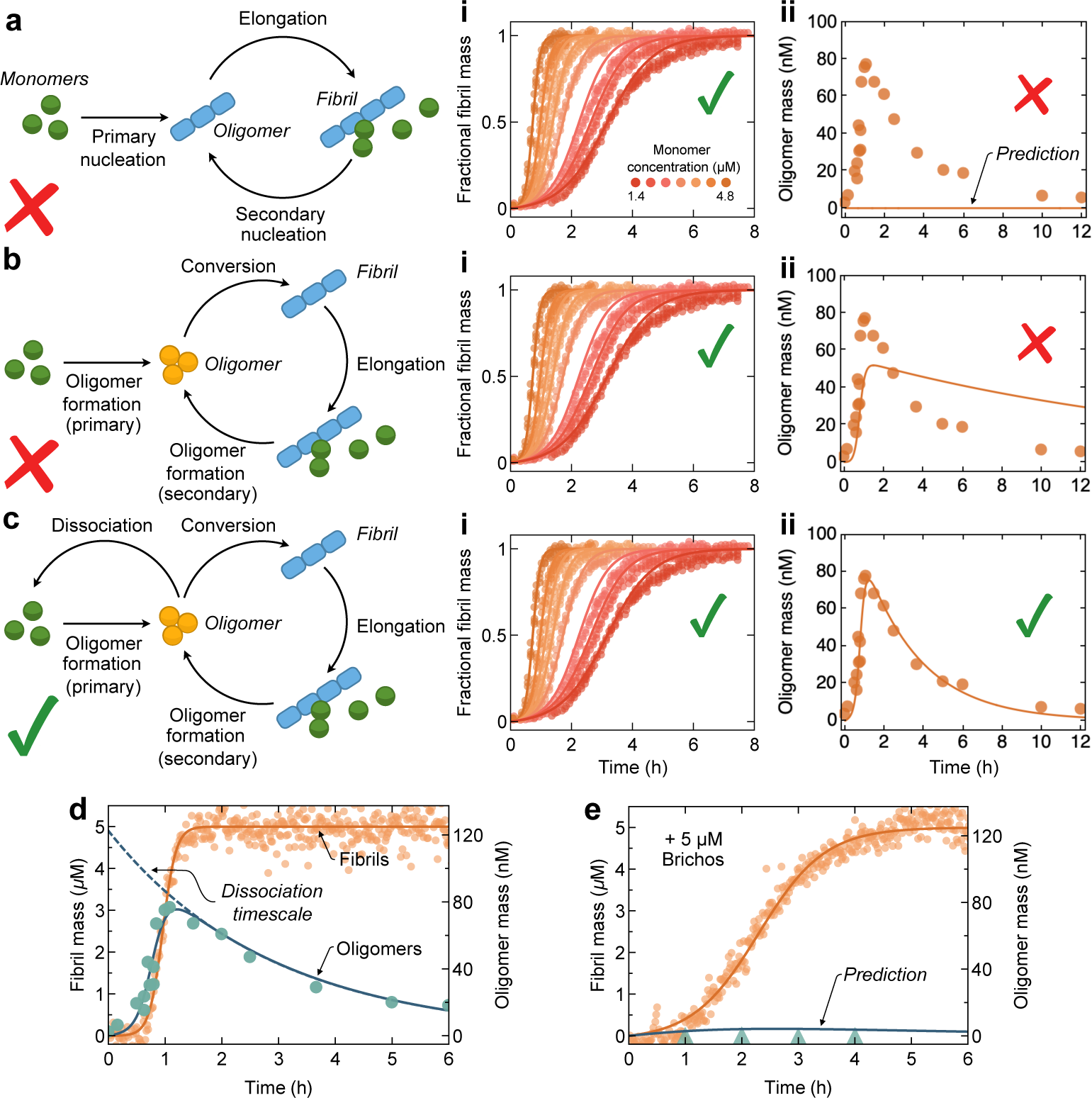
Kinetic analysis of A*β*42 oligomer populations elucidates the molecular pathways of their dynamics during amyloid aggregation. **(a)-(c)** Experimental measurements of (i) fibril formation at varying initial concentrations of A*β*42 (from (16)) and (ii) time evolution of the concentration of oligomers recorded starting from 5*µ*M A*β*42, and best fits (solid lines) to the integrated rate laws corresponding to different mechanistic scenarios for A*β*42 oligomer dynamics (Supplementary Secs. 3-5): **(a)** one-step nucleation producing elongation-competent oligomers, **(b)** two-step nucleation via oligomer conversion to growth-competent fibrils, and **(c)** two-step nucleation via conversion of unstable oligomers. See Supplementary Sec. 6 and Supplementary Table 1 for a detailed description of the fitting procedure and a list of fitting parameters. **(d)-(e)** Experimental measurements of fibril and oligomer kinetics at 5*µ*M A*β*42 in the absence **(d)** and presence **(e)** of 5*µ*M of the Brichos chaperone domain from proSP-C to detect the presence of off-pathway oligomers, i.e. oligomers that do not appreciably contribute to the reactive flux towards fibrils on the time scale of the experiment. Fibril mass measurements were fitted to the analytical expression for the aggregation time course (Supplementary Eq. 25) to determine how the overall rates constants for primary and secondary nucleation are affected by Brichos (Supplementary Sec. 6.4). The rate parameters determined in this way were then used to predict successfully the effect of Brichos on the oligomer concentration, without introducing any additional fitting parameters (Supplementary Eq. 28); this shows that suppressing oligomer formation on fibril surfaces affects equally the reactive fluxes towards oligomers and fibrils, indicating that the majority of oligomers is on-pathway to fibrils.

We next sought to answer the questions of (i) how fast is the conversion rate of oligomers to fibrils and (ii) what fraction of oligomers converts into fibrils or, by contrast, dissociates into monomers without giving rise to new fibrils. To address these questions, we first developed a kinetic model in which oligomeric species undergo a structural conversion before growing into mature fibrils, but cannot dissociate to monomers at a significant rate over the timescale of the experiment (Fig. 2b and Supplementary Sec. 5). While this model is consistent with the fibril mass data (Fig. 2b,i), it cannot reproduce the observed oligomer population dynamics (Fig. 2b,ii). Indeed, in this model the conversion rate controls both the maximum oligomer concentration and the rate of oligomer depletion after this maximum concentration; no value for the conversion rate can simultaneously capture the experimentally observed oligomer peak concentration and the timescale for oligomer depletion. In order to identify the missing element in our analysis we thus introduced in our model an oligomer dissociation process that competes with the conversion into fibrils (Fig. 2c). The inclusion of such a dissociation step in the reaction network does not alter the quality of the fit of the fibril mass data (Fig. 2c,i) but allows the experimental oligomer data to be fitted successfully in this manner (Fig. 2c,ii).

Overall, our oligomer data suggest a two-step mechanism for fibril nucleation involving oligomers as a necessary intermediate step in the formation of fibrils. This mechanism is analogous to the nucleation of crystals in solution, where a liquid state serves as a precursor to the crystal phase.^26,27^ In this two-step nucleation process, a conversion step from oligomers to fibrillar aggregates competes with dissociation of oligomers back to monomers (Fig. 2c). Of these steps, oligomer conversion is on average the slowest under the conditions probed in our experiments. The explicit estimates (Supplementary Table 1) for the (ensemble averaged) rates of conversion (9 × 10^*−*6^ s^*−*1^) and dissociation (9 × 10^*−*5^ s^*−*1^) from this analysis make quantitative predictions, e.g. for the average lifetime of oligomers of about 170 min at 5*µ*M A*β*42. For A*β*42 we thus find the near absence of oligomeric species that are long-lived compared to the timescale of aggregation; this finding could be crucial for determining the extent of spatial propagation of oligomers in living systems from their point of formation on amyloid deposits and plaques.^28^ Equally importantly, the ratio of the conversion and dissociation rates gives the fraction of oligomers that convert successfully to fibril nuclei and eventually transform into mature amyloid fibrils. Strikingly, even though the oligomers are the key source of fibrils, we find that less than 10% of oligomers successfully convert to fibrillar species, whereas the remaining 90% of oligomers dissociate back to the monomeric form.

The next fundamental question is whether the observed oligomers constitute on-pathway species in the process of assembly into fibrils, or are off-pathway structures unable to contribute directly to this process, for example because they cannot convert into fibrils on the timescale of aggregation. To distinguish between the two possibilities, we introduced into the aggregation reaction solution 5*µ*M of the Brichos chaperone domain, which has previously been shown to suppress secondary nucleation very selectively by binding to fibril surfaces, ^29^ and compared its effects both on the population of oligomers and on the reactive flux towards fibrils (Figs. 2d,e). If most oligomers are on-pathway, both processes will be affected equally, whereas if the majority of oligomers is off-pathway, the oligomer concentration and fibril formation rate may be affected in a different manner. The data in the presence of Brichos show that the reactive fluxes towards fibrils and oligomers are affected equally by Brichos, consistent with the majority of the measured oligomeric populations being on-pathway species. A key finding of this work is therefore that the population of oligomers is an ensemble of species interconverting on timescales comparable to aggregation and, hence the fraction of oligomers converting from non-fibrillar forms to growth competent-fibrillar oligomers is determined by the average relative rates of their conversion and dissociation. We also note that *in vivo* some off-pathway oligomers may form from on-pathway ones by being stabilised through the interaction with other cellular components, such as for example metabolites or other proteins.

We then varied the concentration of monomeric A*β*42 in the initial solution to test the ability of our model to capture the observed behaviour and to estimate the reaction orders of the various microscopic processes involved in determining the oligomer populations. These reaction orders coarse grain the monomer concentration dependence of these multistep processes, and contain key information about the rate-determining features of the free energy landscape of amyloid nucleation (Supplementary Sec. 5.1). In particular, we find that the time courses of the oligomer concentrations recorded from solutions with initial monomer concentrations of 2.5*µ*M, 5*µ*M, and 10*µ*M A*β*42 can be described by our model using the same choice of the rate parameters for all 3 datasets, and capturing the dependence on the monomer concentration by means of the reaction orders of the different processes (Fig. 3a). We find that the rate of oligomer conversion displays a marked dependence on monomer concentration with reaction order *n*_conv_ = 2.7, while the oligomer formation step shows a lower reaction order *n*_oligo,2_ = 0.9; these numbers yield an overall reaction order for two-step nucleation of *γ* = (*n*_conv_ + *n*_oligo,2_ + 1)/3 ≃ 1.5, which is equal to the value previously obtained from bulk kinetics assuming single-step nucleation^16^ (Fig. 3c, Supplementary Fig. 17, Supplementary Sec. 5.3, Supplementary Eqs. 24 and 35). Thus, compared to our previous work on secondary nucleation of A*β*42,^16^ with direct measurements and analysis of oligomer populations, we are now able to decompose fibril self-replication into a series of elementary steps, including oligomer formation, conversion, dissociation, and growth, and to quantify the relative importance of each one of these steps. We find that the rate of oligomer depletion is approximatively independent of monomer concentration, while in contrast the fraction of successful oligomer conversion events increases with increasing total protein concentration (Fig. 3b). Overall, these findings suggest that oligomer dissociation is ‘spontaneous’, i.e. independent of oligomer-monomer or oligomer-oligomer interactions, while oligomer conversion involves additional interactions with monomers. This conversion step may occur in solution or in contact with the fibril surface.^24^ Indeed, the latter scenario might explain the high structural specificity of the process. ^30^

**Figure 3:**
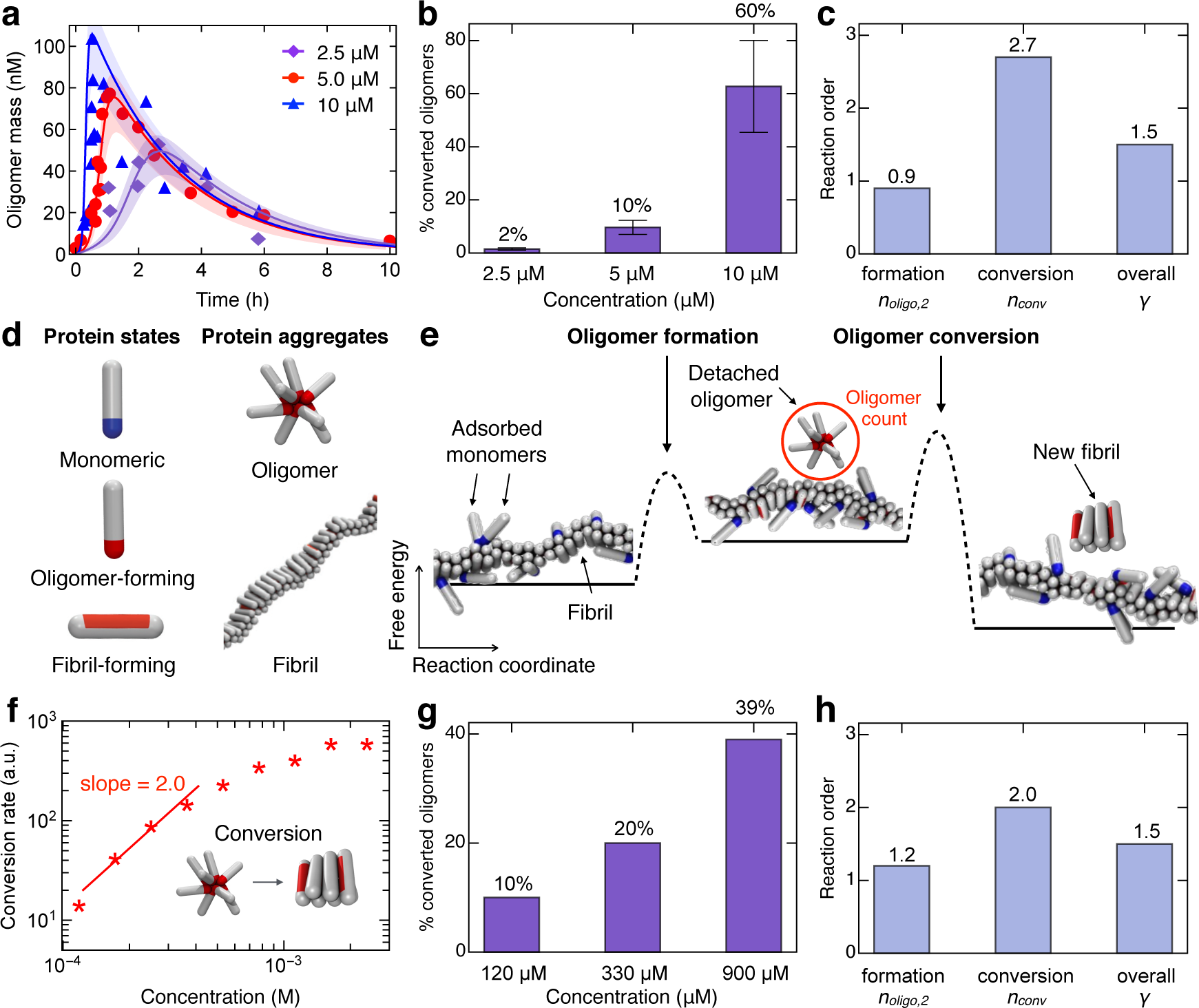
Concentration dependence of the molecular pathways of A*β*42 oligomer dynamics. **(a)-(c)** Experimental measurements of the time evolution of oligomeric populations at varying concentrations of A*β*42 reveal the concentration dependence of oligomer conversion. **(a)** Global fit of experimental oligomer concentration data for 2.5, 5 and 10*µ*M A*β*42 to the integrated rate law corresponding to the model shown in Fig. 2c. Shaded areas indicate 68% confidence bands. See Supplementary Sec. 6 and Supplementary Tables 1 and 2 for a list of fitting parameters. **(b)** Concentration dependence of the fractional contribution of unconverted oligomers towards the reactive flux to mature fibrils. Error bars indicate standard deviation. **(c)** Extracted reaction orders for oligomer formation, oligomer conversion, and overall two-step secondary nucleation. **(d)-(h)** Computer simulation model of A*β*42 aggregation probes concentration dependence of oligomer conversion. **(d)** Possible protein and aggregate states in the computer model. **(e)** Mechanism of secondary nucleation in the computer simulations: monomers adsorb onto the fibril surface, and detach as oligomers, which then convert into fibrils in solution at a later time. However, based on the analysis of our experimental data, we cannot exclude the possibility that structural conversion and dissociation of A*β*42 oligomers occur in contact with, or close, to the fibril surface. **(f)** Rate of conversion of detached oligomers at varying monomer concentrations. **(g)** The fraction of converted oligomers in the total oligomer population at 3 different monomer concentrations. **(h)** Reaction orders for oligomer formation, oligomer conversion and overall two-step secondary nucleation as measured in the simulations.

To provide a structural interpretation of these results, we performed computer simulations using a coarse-grained model of amyloid formation,^31^ which enables us to calculate experimental observables, such as the reaction orders for oligomer formation and conversion, while retaining molecular-level resolution (Figs. 3d-e, Supplementary Sec. 2, Supplementary Fig. 3). In this model, protein monomers are described as single rod-like particles that can interconvert between three states (Fig. 3d): (i) a monomeric state, (ii) an oligomer-forming state, and (iii) a fibril-forming state. The monomeric state represents a disordered monomer in solution, which can adsorb onto the surface of a fibril. The oligomer forming state represents an intermediate state; oligomers formed of particles in this intermediate state can detach from the parent fibril but have not yet converted into a new fibril. Finally, the fibril-forming state represents a *β*-sheet rich state with the ability to form strong lateral interactions. A protein species in its monomeric and oligomer-forming states interacts with particles of the same kind via its ends, that possess an attractive tip. A particle in the fibril-forming state interacts via its sides, that possess an attractive side-patch. This situation mimics directional interactions, such as hydrogen bonding, and drives the formation of fibrillar aggregates. The interaction between two proteins in the fibril-forming state is by far the strongest interaction in the system (Supplementary Fig. 3), and once formed, the growth reaction is effectively irreversible. Every conversion from the monomeric to the fibril-forming state is slow and is thermodynamically penalised by a change in the excess chemical potential, to reflect the fact that amyloidogenic proteins and peptides, such as A*β*, are not typically found in a *β*-sheet conformation in solution.^32,33^ Hence the fibril-forming state is energetically unfavourable. However, as particles in this state interact strongly with other particles of the same kind, the interplay of the two competing energy terms gives rise in the simulations to the nucleation barrier for fibril formation.

We observe in the simulations that oligomers produced via secondary nucleation persist for a significant amount of time in solution before converting to fibrils. Hence, both oligomer formation and conversion are slow steps in the reaction (Supplementary Sec. 2). Moreover, most of the oligomers dissociate back to monomers, and multiple oligomers typically form and dissociate before one successful conversion event into a fibril occurs (Supplementary Movie 1). We can therefore conclude that the fibril surface serves as an oligomer breeding “factory”.^24^ This results in the free energy landscape sketched in Fig. 3e, which involves an initial oligomer formation step followed by a large barrier for oligomer conversion. Note the composite nature of the reaction coordinate, which involves two slow degrees of freedom: aggregate size and structure (*β*-sheet content, see Supplementary Sec. 2.1 and Supplementary Fig. 4 for a discussion). The reaction rates and scaling exponents emerge solely from the molecular ingredients and their interactions (Supplementary Sec. 2.1.3).

As observed in the experiments, oligomer conversion is significantly accelerated at higher monomer concentrations (Fig. 3f), with a high reaction order for oligomer conversion (*n*_conv_ = 2.0) and a low reaction order for oligomer formation (*n*_oligo,2_ = 1.2). These simulations not only reproduce the observed experimental behaviour, but also allow interpretation of the underlying molecular behaviour. For example, we find that larger oligomers have a lower free energy barrier for conversion than smaller ones, rendering the rate of conversion highly dependent on monomer concentration. At higher monomer concentrations, oligomers are not only more numerous but also larger (Supplementary Figs. 5-6), giving rise to faster effective conversion and, hence, to faster overall fibril self-replication. We also find that oligomers can grow in solution, since the size of the oligomers detached from the fibril surfaces is smaller than the average size of converting oligomers (Supplementary Fig. 6). Figure 3g depicts how the fraction of converted oligomers changes with monomer concentration, while the average size of the converting oligomer is depicted in the Supplementary Information. According to our theory, the overall scaling exponent for two-step secondary nucleation can be calculated as *γ* = (*n*_oligo,2_ + *n*_conv_ + 1)/3 ≃ 1.4, which is in excellent agreement with the value measured in the simulations, *γ* = 1.5 (Fig. 3h, Supplementary Eq. 24), as well as in the experiments^16^ thus providing strong support for the mechanistic picture defined in this study.

Finally, we sought to understand how the results for the A*β*42 peptide are applicable to other systems, such as oligomer dynamics of the length variant A*β*40. Using MS (Supplementary Sec. 1.7), we measured the time evolution of the concentration of A*β*40 oligomers starting from a solution with a peptide concentration of 10*µ*M (Supplementary Fig. 8). We then analysed these data using our theoretical framework to determine the rate of oligomer conversion and dissociation for A*β*40 and compare them with those determined for A*β*42 oligomers (Supplementary Table 2). Interestingly, we find that A*β*40 oligomers have a similar rate of conversion (1 × 10^*−*6^ s^*−*1^) compared to A*β*42 oligomers at the same monomer concentration (6 × 10^*−*5^ s^*−*1^), but note that A*β*40 oligomers are somewhat larger than A*β*42 oligomers as they elute earlier from the SEC column (Supplementary Fig. 2). Moreover, from the rate of oligomer dissociation (1 × 10^*−*4^ s^*−*1^) we find that about 1% of the A*β*40 oligomers successfully convert into fibrils, a similar result to A*β*42. Thus, for A*β*40 we arrive to the same conclusion as for A*β*42 that the vast majority of oligomers does not form fibrils, but rather dissociates back to monomers.

## Discussion

We have described an experimental and theoretical approach for elucidating the fundamental molecular pathways driving the dynamics of oligomers during an ongoing amyloid aggregation reaction. By applying this general approach to A*β*42, we have found that, even though all mature amyloid fibrils must originate from oligomers, the majority of oligomers do not convert into fibrils but predominantly dissociate relatively rapidly into monomeric species before the slower conversion step takes place (Fig. 4). This type of mechanism, which fundamentally couples the accumulation of unconverted oligomers with fibril formation, reveals a non-classical nucleation process for A*β*42 amyloid fibrils. The formation of new fibrils occurs in two steps with oligomers as an obligatory intermediate state. The first step is the generation of oligomers through the interaction of monomers with the surface of existing fibrils. The resulting oligomers are a heterogeneous population of aggregates of different size (Supplementary Figs. 2 and 7), which are structurally distinct from fibrils and undergo structural conversions to fibrillar structures. Under the conditions probed in our experiments, the largest free energy barrier associated with fibril nucleation is oligomer conversion. Since intermediate oligomers are unstable species, slow conversion causes most oligomers to dissociate back to monomers without forming new fibrils. The free energy barrier for oligomer conversion is lower at higher monomer concentrations, which explains the observation that the fraction of oligomers that dissociate without forming new fibrils is lower at higher monomer concentrations. The non-classical nucleation behaviour described here for A*β*42 fibril formation is analogous to the two-step nucleation processes observed in crystallisation, bio-mineralisation and sickle-cell haemoglobin.^26,27,34–39^ Moreover, we have established the absence, in our system, of detectable quantities of persistent off-pathway oligomers that cannot convert to fibrils over the timescale of aggregation, although such species may exist under different experimental conditions or for other, particularly larger, amyloidogenic proteins such as *α*-synuclein.^40^ More generally, our work could be extended to study oligomer dynamics in peptide mixtures; in the presence of additional inhibitors,^41^ these experiments could inform upon the role of off-pathway oligomers in such systems. The methods and results described in the present work are also likely to provide essential insights into the rational development of precise therapeutic strategies for targeting oligomers formed during pathological aggregation reactions.

**Figure 4:**
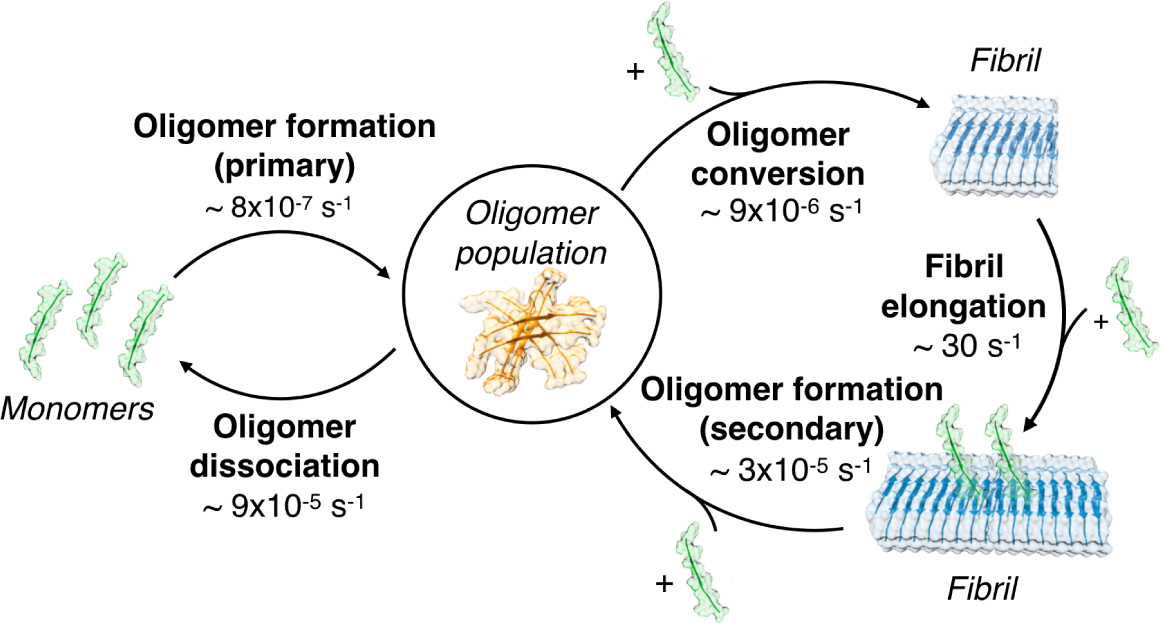
Schematic illustration of the reaction pathways of oligomers during amyloid aggregation and the associated reaction rates determined in this work for A*β*42. Amyloid fibril proliferation occurs through a two-step nucleation mechanism involving oligomer formation followed by oligomer conversion into fibrillar structures. The heterogeneous ensemble of oligomers includes not only converting species but consists mainly of unstable oligomers that can dissociate back to monomers. Oligomers undergo repeated cycles of formation-dissociation before eventually converting into species that are able to grow into new fibrils. The reaction rates are shown here for A*β*42 at a concentration of 5*µ*M (rate constants in Supplementary Sec. 6.3) and are to be interpreted as averages over the heterogeneous ensemble of oligomers. The geometric mean of the rates of oligomer formation, oligomer conversion and fibril elongation (which constitute the autocatalytic cycle of fibril self-replication, Supplementary Sec. 5.3) yields the characteristic rate of amyloid fibril formation (Supplementary Sec. 6.5 and Supplementary Fig. 17).

## Methods

Details of the experimental materials and methods, mathematical modelling, data fitting, and computer simulation model are available in the online version of the paper.

## Acknowledgments

We acknowledge support from Peterhouse, Cambridge (TCTM), the Swiss National Science foundation (TCTM), the Royal Society (AŠ), the Academy of Medical Sciences (AŠ), the UCL Institute for the Physics of Living Systems (SC), Sidney Sussex College, Cambridge (GM), the Wellcome Trust (AŠ, MV, CMD, TPJK), the Schiff Foundation (AJD), the Cambridge Centre for Misfolding Diseases (MV, CMD, TPJK), the BBSRC (CMD, TPJK), the Frances and Augustus Newman foundation (TPJK), the Swedish Research Council (SL) and the ERC grant MAMBA (SL, agreement n° 340890). The research leading to these results has received funding from the European Research Council under the European Union’s Seventh Framework Programme (FP7/2007-2013) through the ERC grant PhysProt (agreement n° 337969).

## Data availability statement

The authors confirm that all data generated and analysed during this study are included in this published article (and its supplementary information files).

## Author contributions

All authors were involved in the design of the study; TCTM developed the theoretical model and performed the kinetic analysis; SL and KB performed the experiments; AS and SC performed computer simulations; all authors participated in interpreting the results and writing the paper.

