## Supplementary Information for "Dynamics of oligomer populations formed during the aggregation of Alzheimer’s A*β*42 peptide"

7 Supplementary Figures 17

8 Supplementary Tables 36

9 Supplementary Movie 37

### 1. Materials and Methods

**1.1. Expression of $^3$H-labelled A$\beta$(M1-42) peptide.** Recombinant human A$\beta$(M1-42) with the sequence MDAEFRHDSGYEVHHQKLVFFAEDVGSNKGAIIGLMVGGVVIA was expressed from a PetSac (derivative of Pet3a) plasmid containing a synthetic gene with *E. coli* optimised codons (see Ref. (1)). The plasmid was transformed into $Ca^{2+}$-competent cells of *E. coli* strain BL21 DE3 PLysS star using heat shock, spread on LB (10 g/L Bacto trypton, 5g/L Bacto yeast extract, 10 g/L NaCl) agar plates (15 g/L Bacto agar) with 50 mg/L ampicillin and 30 mg/L chloramphenicol and grown for 14 h at 37°C. Four 250 mL baffled flasks with 50 mL LB medium with 50 mg/L ampicillin and 30 mg/L chloramphenicol were inoculated with separate well-isolated colonies and grown over night at 37°C and 125 rpm shaking. 3 mL from each overnight culture was transferred to a separate 2,500 mL baffled flask with 500 mL M9 minimal medium (42 mM $Na_2HPO_4$, 22 mM $KH_2PO_4$, 18 mM $NH_4Cl$, 2 g/L glucose, 8.6 mM NaCl, 1 mM $MgCl_2$, 100 $\mu$M $CaCl_2$, 18 $\mu$M $FeCl_3$, 1 $\mu$g/mL vitamin B1, 50 mg/L ampicillin and 30 mg/L chloramphenicol, prepared in Lund tap water). The four M9 cultures were grown at 37°C at continuous shaking at 125 rpm. $^3$H-labelled glucose (with $^3$H in positions 5 and 6 from American Radiolabeled Chemicals Inc., Saint Louis, MO, USA, ART 0312 5mCi) was added to the medium when $OD_{600}$ reached 0.6, and expression was induced 15 minutes later by the addition of 0.4 mM Isopropyl $\beta$-D-1-thiogalactopyranoside (IPTG). Cells were grown for an additional 5 hours and harvested by centrifugation for 10 minutes at 6000 rpm in a JA-8.1000 rotor.

Note that the added $^3$H-glucose was a tracer, and its total fraction was 0.4% of all glucose in the culture. With only a third of the positions labelled we obtain at maximum 0.13% of the carbon-bonded protons replaced with tritium. At all exchangeable positions that can hydrogen bond we have 100% normal hydrogen since the expression and purification was performed in normal water.

**1.2. Expression of $^{15}$N-labelled A$\beta$(M1-42) peptide.** The expression of $^{15}$N-labelled peptide was performed above except that no tritium-labelled glucose was added and $^{15}NH_4Cl_2$ was the only nitrogen source.

**1.3. Expression of unlabelled A$\beta$(M1-42) peptide.** The expression of unlabelled A$\beta$(M1-42) peptide was performed above except that M9 medium was replaced with LB medium with 50 mg/L ampicillin and 30 mg/L chloramphenicol.

**1.4. Expression of unlabelled A$\beta$(M1-40) peptide.** The expression of unlabelled A$\beta$(M1-40) peptide was performed above using a PetSac plasmid with the gene for A$\beta$(M1-40) (see Ref. (1)) and LB medium with 50 mg/L ampicillin and 30 mg/L chloramphenicol.

**1.5. Purification of $^3$H-labelled A$\beta$(M1-42) and A$\beta$(M1-40) peptides.** The peptides were purified from the cell pellet essentially as described (see Ref. (1)) by three times repeated sonication in 80 mL 10 mM Tris/HCl, 1 mM EDTA, pH 7.5, solubilization in 100 mL 10 mM Tris/HCl, 1 mM EDTA, pH 8.5 (buffer A) with 8 M urea, dilution with 300 mL buffer A, application to 50 mL DEAE cellulose ion exchange resin, washing the resin with 100 mL buffer A and elution of the peptide with 8×50 mL buffer A + 50 mM NaCl. All ion exchange steps were performed in batch format. The eluted peptide was lyophylized, dissolved in 6 M GuHCl, 20 mM sodium phosphate buffer, pH 8.5 and subjected to two rounds of size exclusion chromatography on a 26×600 mm Superdex 75 column in 20 mM sodium phosphate buffer, 0.2 mM EDTA, pH 8.5 (buffer B) with intervening lyophylization and dissolution in 6 M GuHCl, 20 mM sodium phosphate buffer, pH 8.5. The resulting monomer was lyophylized as multiple identical aliquots.

**1.6. Aggregation kinetics, oligomer isolation and quantification.** For each kinetic experiment, one $^3$H-A$\beta$(M1-42) monomer aliquot from above was dissolved in 1 mL 6 M GuHCl, 20 mM sodium phosphate buffer, pH 8.5 and monomer again isolated by size exclusion chromatography on a 10×300 mm Superdex 75 column in 20 mM sodium phosphate buffer, 0.2 mM EDTA, pH 8.0. The centre of the monomer peak was collected in a low-binding tube (Axygen) on ice, diluted to the desired working concentration (10, 5 or 2.5 $\mu$M) and supplemented with 5 $\mu$M Thioflavin T (ThT). Multiple 100 $\mu$l samples were placed in individual wells of a 96 well half-area low-binding plate (PEG-ylated polystyrene, Corning 3881) and the aggregation reaction initiated by placing the plate in a plate reader at 37°C quiescent. Fibril formation was monitored by recording ThT fluorescence through the bottom of the plate (excitation 440 nm, emission 480 nm). For each time point, $\Delta t$, the plate was stopped, sample from twelve wells combined, centrifuged for 2 min at 18000 rpm in a 2 mL low-binding tube (Axygen) and the top 1mL of the supernatant injected on a 10×300 mm Superdex 75 column in 20 mM sodium phosphate buffer, 0.2 mM EDTA, pH 8.0. The fractions between the void and the monomer peak were collected, lyophilised, dissolved in 250 $\mu$l $H_2O$ and mixed with 2.25 mL Ultima Gold scintillator liquid (Perkin Elmer, Waltham, MA) and the radio decays counted using a Beckman LG6000IC liquid scintillator counter. A standard curve in the range 0-200 nM was prepared from the monomer solution.

Note that the total time for the oligomer isolation process (from collecting the samples until starting the SEC run) is 3 minutes. Oligomer dissociation is negligible over this time scale: the estimated rate of oligomer dissociation $9.7 \times 10^{-5}$ s$^{-1}$ (see Supplementary Sec. 6.3) predicts in fact that less than 2% of oligomers would dissociate during the isolation process. Moreover, the SEC separation process takes 8-13 minutes (oligomers are collected as the fraction eluting between 8 and 13 min after start) but only oligomers that dissociate very early during the SEC procedure will be retarded beyond 13 minutes. Therefore, most oligomers that dissociate during the SEC run will be retained in the collected oligomer fraction.

**1.7. Oligomer quantification by mass spectrometry.** A$\beta$(M1-42) monomer was isolated by size exclusion chromatography as above in 20 mM sodium phosphate buffer, 0.2 mM EDTA, pH 8.0, diluted to 5 $\mu$M and supplemented with 5 $\mu$M ThT. A$\beta$(M1-40) monomer was isolated by size exclusion chromatography as above in 20 mM sodium phosphate buffer, 0.2 mM EDTA, pH 7.4, diluted to 10 $\mu$M and supplemented with 20 $\mu$M ThT. The aggregation reaction was followed as above and samples from twelve wells were combined at each time point, centrifuged for 2 min at 18000 rpm in a 2 mL low-binding tube (Axygen) and the top 1 mL of the supernatant injected on a 10$\times$300 mm Superdex 75 column in 20 mM ammonium acetate, pH 8.5. Fractions between the void and the monomer peak were collected, and lyophylised. Each fraction was dissolved in 20 $\mu$L $H_2O$, supplemented with 10 pmol $^{15}$N-A$\beta$(M1-42), subjected to digestion with AspN protease overnight. The sample was then purified by HPLC using a C18 column and spotted on a MALDI-plate, followed by application of $\alpha$-Cyano-4-hydroxycinnamic acid matrix solution and mass spectrometric analysis using an Autoflex Speed MALDI TOF/TOF system (Bruker Daltonics). The relative intensity of the peaks at 1906 m/z ($^{14}$N-A$\beta$7-22, DSGYEVHHQKLVFFAE) and 1928 m/z ($^{15}$N-A$\beta$7-22 isotope standard), was used to estimate the oligomer concentration in each fraction (Supplementary Fig. 2).

### 2. Computer Simulations

We used the coarse-grained computer simulation model for secondary amyloid nucleation developed in Ref. (2), based on our previous work published in Ref. (3). This model is simple enough to allow us to explore possible mechanistic scenarios and their kinetics, depending on the design of the inter-protein interactions in the system. In this model, the protein is described by a single rod-like particle which can interconvert between three states: the monomeric state ('s') in which it can adsorb onto the surface of a fibril, the oligomer-forming intermediate state ('i') that has detached from the parent fibril but not converted into a new fibril yet, and a fibril-forming state ('$\beta$'). A protein in its monomeric and oligomer-forming states interacts with its own kind via its attractive tip, while a particle in the fibril-forming state interacts with its attractive side-patch, mimicking directional interactions, such as hydrogen bonding, and drives the formation of fibrillar aggregates. The interaction between two proteins in the fibril-forming states is by far the strongest interaction in the system (Supplementary Fig. 3), and once formed, the fibrils are effectively irreversible. Every conversion from the monomeric state to the fibril-forming state is penalised with a change in the excess chemical potential of magnitude $\Delta\mu = 20\,kT$, where $kT$ is the thermal energy. This reflects the fact that amyloidogenic proteins such as A$\beta$ are typically not found in the $\beta$-sheet prone conformation in solution (4, 5). Hence the fibril-forming state is energetically unfavourable, but a particle in this state interacts strongly with its own kind; it is this interplay of the two competing energy terms which gives rise to the nucleation barrier for fibril formation.

Monomer adsorption onto the fibril is energetically favourable (Supplementary Fig. 3), and monomers are able to interact on the fibril surface and form oligomers made of proteins in the monomeric state. Since the protein in the oligomer-forming state interacts with the fibril only weakly (Supplementary Fig. 3), oligomer detachment is favourable only for large enough oligomers, where the loss of monomer-fibril interactions is overcome by the gain by the interaction between proteins in the oligomer-forming state. In our previous work (2), the conversion from the monomeric state to the oligomer-forming state, as well as the conversion from the oligomer-forming state to the fibril forming state were penalised by $0.5\Delta\mu$. This created a large barrier for the formation of oligomers and their detachment, but once oligomers were formed, they quickly converted into a fibril (with $n_{\mathrm{conv}} = 0.5$). To approach the situation observed in experiments presented in this paper, we adjusted the interaction parameters in our model (Supplementary Fig. 3), by lowering the interaction strength between proteins in the oligomer-forming state, not penalising the conversion from the monomeric state to the oligomer-forming state, while penalising the conversion from the oligomer-forming state to the fibril-forming state by the full $\Delta\mu$. This effectively caused the oligomer-forming protein state to be closer to the native state than to the fibril-forming state in the free energy landscape. As a consequence, the oligomers produced via the secondary pathway experience a significant barrier for the conversion into the fibril, rendering this step rate-determining, as reflected by the large value of the reaction order.

**2.1. Insights into nucleation mechanism of amyloid fibrils.** Our analysis of A$\beta$42 oligomers provides insights into the nucleation of amyloid fibrils, suggesting a non-classical nucleation mechanism through non-fibrillar oligomers. This nucleation mechanism is analogous to the two-step nucleation mechanism observed for crystallization and biomineralization (6, 7).

***2.1.1. Classical nucleation theory.*** Classical nucleation theory (CNT) assumes that the only relevant degree of freedom in the system is the physical size of aggregates (Supplementary Fig. 4(a)). There is a free energy barrier for nucleation only when large aggregates are more stable compared to the monomer phase. In this case, the reaction order of nucleation (or critical nucleus size) is necessarily related to the physical size of the nucleus. This result, which is generally known as the nucleation theorem (8), is a direct consequence of the fact that CNT has only one relevant degree of freedom.

***2.1.2. Non-classical nucleation of amyloid fibrils.*** Unlike CNT, in our computer simulations, nucleation of new fibrils proceeds through disordered micellar-like oligomers, whose subunits are changing conformation during nucleation. This is a consequence of the fact that proteins in the final, fibrillar state acquire a $\beta$-sheet conformation, which is very different from the conformation of monomers in solution. This conformational change from monomeric to $\beta$-sheet form is energetically unfavourable and slow. This situation leads to a nucleation process involving the fraction of $\beta$-sheet conformation within the cluster as an additional slow degree of freedom in addition to aggregate size (Supplementary Fig. 4(b)). This corresponds to a two-step nucleation process, as introduced by Vekilov and coworkers (6) and discussed in the context of amyloid nucleation in Refs. (3, 9–11). Computer simulations allow us to relate these findings to a microscopic picture. In the simulations, we observe that larger oligomers have a lower free energy barrier for conversion than smaller ones, rendering the rate of conversion dramatically dependent on the monomer concentration. At higher monomer concentrations the oligomers not only are more numerous (Supplementary Fig. 5) but also larger (Supplementary Fig. 6), giving rise to faster conversion and, hence, overall fibril self-replication. Consequently, the fraction of converted oligomers increases with monomer concentration. We find that the average oligomer size in the system increases with increasing monomer concentration. We also find that the average size of the converting oligomer increases with increasing monomer concentration (Supplementary Fig. 6). The average size of converting oligomers is larger than the ensemble average over the entire oligomer population (Supplementary Fig. 6). This is consistent with the idea that only the largest oligomers from the size distribution can convert. Note that the reaction order for conversion measured in the simulations ($n_{\mathrm{conv}} = 2$) is not equal to the actual size of converting oligomers.

***2.1.3. Link to kinetic equations.*** The situation that emerges from our computer simulations corresponds to that described by our kinetic equations Eq. (11) (see Supplementary Sec. 5.1). Once detached, the oligomers nucleated on the surface of the fibril seed spend a significant time in solution before converting into a fibril. During this period, oligomers have a significant chance to dissolve back to monomers. Thus, the reaction flux towards fibrils is determined by a competition between oligomer conversion and dissociation (last two terms on the right-hand side of Eq. (11a) and right-hand side of Eq. (11b)). The oligomer population constitutes of oligomers of different sizes (Supplementary Fig. 7). We find that oligomers can grow in solution, since the size of detaching oligomers is smaller than the average oligomer size or the size of the nucleating oligomer (Supplementary Fig. 6). We also find that oligomer conversion is significantly faster for larger oligomers. This causes the rate of oligomer conversion to display a marked dependence on monomer concentration, a feature which is captured by the reaction order $n_{\mathrm{conv}}$ for conversion. As discussed here below, $n_{\mathrm{conv}}$ is determined by the size of the portion of the oligomer that converts first - the number of proteins that are in contact during the first conversion step - which is not necessarily equal to the size of the whole oligomer.

It is important to note that the terms in the kinetic equations Eq. (11) are not explicitly implemented in the simulations. Our simulations consider only the molecular ingredients and their interactions described in Supplementary Sec. 2. Solely from these molecular ingredients we recover the terms from our kinetic model. This is indeed the power of such coarse-grained simulation models: they allow us to simulate molecular details and reach experimentally relevant time-scale, therefore effectively bridging molecular and continuum scales (12).

### 3. Master equation approach to oligomer dynamics

In this section, we describe in detail a master equation approach to study the dynamics of oligomer populations formed during amyloid fibril formation. The first step of this approach consists in identifying the most general set of physically meaningful microscopic events through which oligomeric intermediates are populated and depleted (see Supplementary Sec. 3.1). We then account for these various possible elementary events by means of master equations (see Supplementary Secs. 4.1 and 5.1). A master equation is a set of coupled differential equations that explicitly quantify the population balance of each of the species relevant to the aggregation reaction in terms reaction fluxes (13–17). In order to perform a quantitative comparison between experimental data and the model predictions, we solve the master equations using self-consistent approaches to yield integrated rate laws (see Ref. (18) for an introduction to this mathematical technique). In this manner, we can overcome the non-linear nature of the master equations and derive explicit analytical solutions to the dynamics of the fibril mass concentration (see Supplementary Secs. 4.2 and 5.4) as well as the concentration of oligomers (see Supplementary Secs. 4.3 and 5.5). Such integrated rate laws can be used to determine the relative importance of the different rate parameters from experimental data.

**3.1. Microscopic mechanisms of oligomer dynamics.** We now consider in detail all possible elementary aggregation events that, in a system consisting of monomers and aggregates, are responsible either for increasing or for decreasing the population of oligomers. We distinguish two main classes of microscopic processes contributing to oligomer dynamics:

(1) Oligomer formation events are responsible for populating oligomeric intermediates. In a system containing monomers and aggregates, oligomers could be formed in a manner that depends either on the free monomers alone, on the fibrils alone or on any combination of monomer and fibril populations. In the most general scenario, we thus distinguish between primary (i.e. fibril-independent) oligomer formation events, where one or more monomers come together to form new oligomers, and secondary (i.e. fibril-dependent) oligomer formation events, where one or more monomers interact with the surface of existing fibrils to generate new oligomers. Any possible dependence of primary and secondary oligomer formation steps on the free monomer concentration is captured by means of reaction orders.

(2) Oligomer depletion events are responsible for decreasing the population of oligomers. Very generally, reactive fluxes away from oligomeric intermediates can be directed either (i) towards mature fibrils or (ii) towards the monomers and hence involve dissociation events. If oligomers are short fibrils (i.e. elongation-competent species that are capable of growing by sequestering free monomers, see Fig. 2a of the main text and mechanism 1), then any decrease of the population of oligomers in favor of mature fibrils is due to the elongation of oligomers into longer fibrillar species. Instead, if oligomers are distinct from short fibrils (i.e. are not elongation-competent, see Figs. 2b and 2c of the main text and mechanisms 2 and 3), then the reactive flux from oligomer concentrations to mature fibrils must involve conversion events into fibrillar species that are capable of further growth. Note that, since all mature fibrils must originate from oligomers, a reactive flux from oligomers towards mature fibrils must necessarily exist, either in the form of elongation or conversion events. Also note that each one of these oligomer depletion events could depend on the monomer population. Here, we allow for this possibility by assigning to each of these steps a reaction order with respect to the monomer concentration: $n_{\mathrm{conv}}$ for oligomer conversion and $n_d$ for dissociation. In accordance with previous reports (16), we take the reaction order for elongation to be 1 (i.e. the rate of elongation depends linearly on the concentration of free monomer), but note that at monomer concentrations higher than those considered in this study saturation effects in elongation can become important (19). Our approach could be generalized in a straight-forward way to account for this effect.

Taken together, these classes of molecular-level events are the basis of a general description of oligomer formation and cover the possible scenarios where mixed populations of monomers and aggregates contribute to the increase or decrease of oligomeric populations (see list of possible mechanistic scenarios in Fig. 2 of the main text and Supplementary Figs. 9 and 11). In Supplementary Secs. 4 and 5, we develop in detail a master equation formalism to describe each one of these mechanistic scenarios. We then employ self-consistent methods to derive explicit mathematical expressions for the fibril mass and oligomer concentrations which are used in the main text to compare model predictions with the experimental data and hence draw a detailed mechanistic picture of oligomer dynamics during A$\beta$42 amyloid fibril formation. The description of the fitting procedure is given in Supplementary Sec. 6.

#### 3.2. Notes on nomenclature.

***3.2.1. Oligomer nomenclature.*** Note the different meaning of the term “oligomer” in the context of mechanism 1 (Supplementary Sec. 4, Supplementary Fig. 9 and Fig. 2a of the main text) and mechanisms 2 and 3 (Supplementary Sec. 5, Supplementary Fig. 11 and Figs. 2b,c of the main text).

- In mechanism 1 (Supplementary Sec. 4, Supplementary Fig. 9 and Fig. 2a of the main text) the term “oligomer” refers to small fibrillar structures with size up to ca. 20 monomers. These oligomers can thus be thought of as “fibrillar oligomers”; in fact, they are able to elongate in a manner similar to that established for mature amyloid fibrils and thus behave kinetically in the same way as mature fibrils.

- In mechanisms 2 and 3 (Supplementary Sec. 5, Supplementary Fig. 11 and Figs. 2b,c of the main text), the term “oligomer” is used to denote “unconverted oligomers” that, unlike fibrillar structures, are not capable of growing directly into mature fibrils but must instead convert into short fibrils first before being able to undergo elongation processes. For simplicity, we shall use the term "oligomer" only to denote unconverted oligomers and shall refer to converted oligomers as "fibrillar oligomers" or simply as "fibrils". Note that this choice of nomenclature is fully consistent with our experimental measurements of oligomer populations; indeed, the majority component of the measured oligomer populations consists of unconverted oligomers while converted oligomers (fibrillar oligomers) occur at concentrations that are much smaller than those of unconverted oligomers (see Supplementary Fig. 14).

***3.2.2. Oligomer formation vs nucleation.*** Also note the difference between oligomer formation events and nucleation events. Nucleation is defined as the process of formation of new fibrils, while oligomer formation events refer to the process of formation of new oligomers.

- In the case of mechanism 1 (Supplementary Sec. 4, Supplementary Fig. 9 and Fig. 2a of the main text), oligomers are small fibrils. Thus, the primary and secondary oligomer formation steps are in this case (and only in this case) equivalent to the primary and secondary nucleation of new fibrils and the terms "oligomer formation" and "nucleation" can be used interchangeably.

- In mechanisms 2 and 3 (Supplementary Sec. 5, Supplementary Fig. 11 and Figs. 2b,c of the main text), the primary and secondary oligomer formation steps give rise to an heterogeneous mixture of unconverted oligomers that convert to fibrillar structures or dissociate back to monomers. In this case, primary and secondary oligomer formation events are different from the respective nucleation events; oligomer formation is just a sub-step during the nucleation of new fibrils. In this case, primary and secondary nucleation are broken up into their constituent steps of oligomer formation and conversion.

### 4. Mechanism 1

Here, we consider the situation when the oligomers generated through the primary and secondary nucleation pathways are small fibrillar aggregates (Supplementary Fig. 9). For clarity, we shall refer to this scenario as "mechanism 1". Below we describe the master equation that captures the system dynamics in this limit (see Supplementary Sec. 4.1) and present the derivation of integrated rate laws for the aggregate mass (see Supplementary Sec. 4.2) and oligomer concentrations (see Supplementary Sec. 4.3).

**4.1. Master equation.** The time evolution of a system described by mechanism 1 (Supplementary Fig. 9) is captured by the following master equation, as described in previous work (13–17):

$$\begin{aligned}\frac{\partial f(t,j)}{\partial t} &= 2k_+ m(t) f(t,j-1) - 2k_+ m(t) f(t,j) \\ &\quad + k_1 m(t)^{n_1}\,\delta_{j,n_1} + k_2 m(t)^{n_2} M(t)\,\delta_{j,n_2},\end{aligned} \tag{1}$$

where $f(t,j)$ is the time-varying concentration of aggregates of size $j$, $m(t)$ is the monomer concentration, $M(t) = m_{\text{tot}} - m(t)$ is the aggregate mass concentration, $m_{\text{tot}}$ is the (conserved) total concentration of monomers, $k_+$, $k_1$ and $k_2$ are the rate constants for elongation, primary and secondary nucleation, and $n_1$ and $n_2$ are the reaction orders of the primary and secondary nucleation steps with respect to the concentration of monomers. Note that, in this model, primary and secondary nucleation are described as single-step processes. The concentration of oligomers in this case is defined as (17):

$$O(t) = \sum_{j=\min\{n_1,n_2\}}^{j_{\max}} f(t,j),$$

where $j_{\max}$ is the maximal size of oligomers.

**4.2. Solution for aggregate mass and monomer concentrations.** A convenient strategy for dealing with master equations such as Eq. (1) is to focus first on the time evolution of the aggregate number and mass concentrations

$$P(t) = \sum_{j=\min\{n_1,n_2\}}^{\infty} f(t,j), \quad M(t) = \sum_{j=\min\{n_1,n_2\}}^{\infty} j f(t,j). \tag{2}$$

This can be obtained form Eq. (1) through summation over aggregate size $j$, yielding the following set of coupled differential equations (13–17):

$$\frac{dP(t)}{dt} = k_1 m(t)^{n_1} + k_2 m(t)^{n_2} M(t) \tag{3a}$$

$$\frac{dM(t)}{dt} = 2k_+ m(t) P(t) + n_1 k_1 m(t)^{n_1} + n_2 k_2 m(t)^{n_2} M(t) = -\frac{dm(t)}{dt}. \tag{3b}$$

Note that the dominant term contributing to the increase of aggregate mass concentration (or, equivalently, decrease of monomer concentration) is the elongation of existing aggregates at their ends with a rate constant $k_+$ and not the consumption of monomers during primary and secondary nucleation; otherwise, aggregate lengths would be much smaller than observed (several thousands of monomers). Hence, the nucleation terms in the equation for $dM(t)/dt$ are negligible relative to the elongation term, and we arrive at the following simplified equations (13–17):

$$\frac{dP(t)}{dt} = k_1 m(t)^{n_1} + k_2 m(t)^{n_2} M(t) \tag{4a}$$

$$\frac{dM(t)}{dt} = 2k_+ m(t) P(t) = -\frac{dm(t)}{dt}. \tag{4b}$$

An accurate analytical solution to Eq. (4) when $m(0) = m_{\text{tot}}$ can be obtained using self-consistent methods by combining the early- and late-time dynamics of the system into a single expression (13, 17):

$$\frac{M(t)}{m_{\text{tot}}} = 1 - \frac{m(t)}{m_{\text{tot}}} = \frac{\frac{\lambda^2}{2\kappa^2}\left(e^{\kappa t} - 1\right)}{1 + \frac{\lambda^2}{2\kappa^2}\left(e^{\kappa t} - 1\right)}, \tag{5}$$

where

$$\lambda = (2k_+ k_1 m_{\text{tot}}^{n_1})^{1/2}, \quad \kappa = (2k_+ k_2 m_{\text{tot}}^{n_2+1})^{1/2} \tag{6}$$

are effective rates that describe the proliferation of aggregates through primary and secondary nucleation, respectively.

**4.3. Estimating the concentration of oligomers.** We now demonstrate how Eq. (5) can be used to estimate the concentration of oligomers for the scenario described by mechanism 1. To do so, it is useful to first replace in Eq. (1) the discrete aggregate size variable $j$ by a continuous variable $x$ and replace finite differences with derivatives:

$$f(t, j-1) - f(t, j) \simeq -\frac{\partial f(t, x)}{\partial x}.$$

This operation transforms Eq. (1) into the following advection equation

$$\frac{\partial f(t, x)}{\partial t} = -v(t)\frac{\partial f(t, x)}{\partial x} + \left[k_1 m(t)^{n_1} + k_2 m(t)^{n_2} M(t)\right] \delta(x - n_2), \tag{7}$$

where the advection velocity in $x$-space is $v(t) = 2k_+ m(t)$ and for simplicity we set $n_1 = n_2$. The solution to the advection equation Eq. (7) is given by a traveling wave in aggregate size space $f(t, x) = F(x - \tau(t))$, where $\tau(t) = \int_0^t v(s)ds$ quantifies the total amount of monomers depleted until time $t$. The profile function $F$ of the traveling wave solution is determined from the boundary condition at $x = n_2$, which can be obtained by integrating Eq. (7) with respect to $x$ over the interval $[n_2, n_2 + \varepsilon]$ and letting $\varepsilon$ to zero:

$$f(t, n_2) = \frac{k_1 m(t)^{n_1} + k_2 m(t)^{n_2} M(t)}{v(t)}. \tag{8}$$

Thus, in the case when oligomers correspond to the smallest fibrillar structure ($j = n_2$), we obtain the following estimate for the overall oligomer concentration:

$$O(t) = \frac{k_1}{2k_+} m(t)^{n_1 - 1} + \frac{k_2}{2k_+} m(t)^{n_2 - 1} M(t). \tag{9}$$

Interestingly, according to Eq. (9) the time course of the fibrillar oligomer concentration depends only on the aggregate (and monomer) concentration. Thus, the concentration of oligomers over time can be predicted solely on the basis of the kinetic parameters that can be extracted from an analysis of the aggregate mass concentration in bulk (i.e. $k_+$, $k_1$ and $k_2$), without the need of introducing additional fitting parameters. Supplementary Fig. 10 shows the time course of fibril mass concentration and the concentration of fibrillar oligomers from a initially monomeric solution ($m(0) = m_{\text{tot}} = 5\mu$M) predicted using the values for the rate constants $k_+$, $k_1$ and $k_1$ estimated previously in Ref. (16) from fitting the fibril mass data of Fig. 2a,i of the main text to Eq. (5). From this figure it is obvious that model 1 underestimates by at least 5 orders of magnitude the concentration of oligomers (sub-pM range) compared to our experimental measurements (nM range). This fact remains true even if we consider the fact that oligomeric populations measured in our experiments comprise aggregates with size from 2- to 20-mers. In this case, a generous estimate of the predicted oligomer concentration would be to consider 20 times the prediction of $O(t)$. Under these conditions, the peak concentration of oligomers can be estimated from Eq. (9) using these rate parameters as:

$$O_{\max} \leq 20 \times \frac{k_2 m_{\text{tot}}^{n_2}}{2^{n_2+1} k_+}. \tag{10}$$

This equation yields an estimate for $O_{\max}$ of 280 fM. This predicted value is about $3 \times 10^5$ times smaller than the measured oligomer peak concentration of 78 nM (see Fig. 2a,ii of the main text).

### 5. Mechanisms 2 and 3

Since mechanism 1 is not able to explain the experimental time course of oligomer concentrations measured in this study, we now consider the scenario when oligomers generated through primary and secondary nucleation undergo a conversion step into fibrillar species before they can grow into mature fibrils by recruiting monomers at their ends (Supplementary Fig. 11). For clarity, we shall refer to this as "mechanism 2". In "mechanism 3", we allow unconverted oligomers to undergo dissociation events back to monomers in addition to conversion events to fibrillar structures.

**5.1. Kinetic equations.** To model the system dynamics for mechanisms 2 and 3, we first write down the associated differential rate law. This can be done by extending Eq. (4) to account for an additional species (oligomers) that are able to convert to fibrils or dissociate back to monomers without forming new fibrils:

$$\frac{dO(t)}{dt} = k_{\text{oligo},1}m(t)^{n_{\text{oligo},1}} + k_{\text{oligo},2}m(t)^{n_{\text{oligo},2}}M(t) - [k_c m(t)^{n_{\text{conv}}} + k_d]O(t) \quad \text{[11a]}$$

$$\frac{dP(t)}{dt} = k_c m(t)^{n_{\text{conv}}}O(t) \quad \text{[11b]}$$

$$\frac{dM(t)}{dt} = 2k_+ m(t)P(t) = -\frac{dm(t)}{dt}, \quad \text{[11c]}$$

where $M(t)$ and $P(t)$ are the mass and number concentrations of aggregates, $m(t)$ is the concentration of monomers and $O(t)$ is the concentration of (unconverted) oligomers. $k_{\text{oligo},1}$ is the rate constant for the oligomer formation step in primary nucleation, $k_{\text{oligo},2}$ is the rate constant for oligomer formation step in secondary nucleation, $k_c$ is the conversion rate constant, $k_d$ is the rate for oligomer dissociation, $k_+$ is the rate constant for fibril elongation, $n_{\text{oligo},1}$ and $n_{\text{oligo},2}$ are the reaction orders for the primary and secondary oligomer formation steps, $n_{\text{conv}}$ is the reaction order for conversion.

It is important to note that both primary and secondary nucleation of new filaments are non-classical nucleation processes, involving aggregate size as well as structural degrees of freedom (associated with $\beta$-sheet content) as slow reaction coordinates. The rate laws employed in our model equation are a coarse-grained descriptions of these multi-step processes. We coarse grain these complicated multistep processes by means of power laws because we are interested in determining the associated reaction orders. These reaction orders reflect the monomer concentration dependence of these multistep processes and hence contain key information about the rate-determining features of the free energy landscape. Coarse-grained computer simulations (see Supplementary Sec. 2) can help us gaining a molecular-level understanding of the meaning of these reaction orders.

Note that, in general, the rate of dissociation in Eq. (11), $k_d$, could have an intrinsic concentration dependence, which would account for the possibility that oligomer dissociation not being a unimolecular step. In this case, we would write the dissociation rate as $k_d = k'_d m(t)^{n_\text{d}}$, where $k'_d$ is the dissociation rate constant and $n_d$ is the reaction order of dissociation. Concentration-dependent dissociation rates have been measured for different systems, such as DNA-binding proteins; in this example, the concentration dependence emerges since dissociation is a multi-step process involving partial dissociation and exchange with other proteins in solution (21). While our A$\beta$42 oligomer data suggest that oligomer dissociation is a unimolecular reaction in this case, we cannot rule out the possibility that other protein systems exhibit a concentration dependence of oligomer dissociation.

As for Eq. (4), to derive Eq. (11) we have used the fact that fibril elongation is the main generator of aggregate mass, such that nucleation, conversion and dissociation terms can be neglected in front of the elongation term in the equation for $dM(t)/dt$. Finally, we note that the description of mechanism 2 follows directly from Eq. (11) with $k_d = 0$. In the following, we will consider Eq. (11) together with the initial conditions $O(0) = M(0) = P(0) = 0$.

The system of equations Eq. (11) has no explicit exact analytical solution; however, accurate approximate solutions can be obtained using self consistent methods, which have successfully been applied in the past to attack similar problems relating to filament growth (13–18). The general idea of this approach is to rewrite, through formal integration, the dynamic equations Eq. (11) as a fixed-point equation. The resulting fixed-point equation is solved iteratively by repeated application of the associated fixed-point operator to an initial guess for the solution. As the starting point of this iterative procedure we choose the solution to the linearized moment equations Eq. (11), which accurately describes the system dynamics up to a time where the growth of aggregates deviates from an exponential growth and begins to saturate due to depletion of monomers.

**5.2. Linearized kinetic equations – Early stages of aggregation.** We start our analysis of the solution to Eq. (11) by restricting ourselves to the early stages of the aggregation dynamics, when the monomer concentration hardly changes. During these early stages of dynamics, we can thus linearize the right hand side of Eq. (11) by setting $m(t) = m_{\text{tot}}$ throughout, yielding:

$$\frac{dO_0(t)}{dt} = \alpha_1 + \alpha_2 M_0(t) - \rho_e O_0(t) \quad \text{[12a]}$$

$$\frac{dP_0(t)}{dt} = \rho_c O_0(t) \quad \text{[12b]}$$

$$\frac{dM_0(t)}{dt} = \mu P_0(t), \quad \text{[12c]}$$

where the index $_0$ is used to indicate solutions to the linearized equations and $\alpha_1 = k_{\text{oligo},1}m_{\text{tot}}^{n_{\text{oligo},1}}$, $\alpha_2 = k_{\text{oligo},2}m_{\text{tot}}^{n_{\text{oligo},2}}$, $\rho_c = k_c m_{\text{tot}}^{n_{\text{conv}}}$, $\rho_e = k_c m_{\text{tot}}^{n_{\text{conv}}} + k_d$, $\mu = 2k_+ m_{\text{tot}}$. The (early-time) linearized moment equations Eq. (12) can be re-written in matrix form as:

$$\frac{\mathrm{d}}{\mathrm{dt}}\begin{pmatrix} O_0(t) \\ P_0(t) \\ M_0(t) \end{pmatrix} = \begin{pmatrix} -\rho_e & 0 & \alpha_2 \\ \rho_c & 0 & 0 \\ 0 & \mu & 0 \end{pmatrix}\begin{pmatrix} O_0(t) \\ P_0(t) \\ M_0(t) \end{pmatrix} + \begin{pmatrix} \alpha_1 \\ 0 \\ 0 \end{pmatrix}. \quad \text{[13]}$$

The solution for the oligomer concentration is of the form $O_0(t) = C_1 e^{x_1 t} + C_2 e^{x_2 t} + C_3 e^{x_3 t}$, where $x_i$ are the eigenvalues of the matrix in Eq. (13), given by the roots of the following cubic characteristic equation:

$$x^3 + \rho_e x^2 - \alpha_2 \rho_c \mu = 0 \quad \text{[14]}$$

and $C_i$'s are constants of integration given by:

$$C_1 = \frac{\alpha_1(x_3 + x_2 + \rho_e)}{(x_1 - x_2)(x_3 - x_1)},$$
$$C_2 = \frac{\alpha_1(x_3 + x_1 + \rho_e)}{(x_2 - x_1)(x_3 - x_2)}, \quad [15]$$
$$C_3 = \frac{\alpha_1(x_2 + x_1 + \rho_e)}{(x_3 - x_1)(x_2 - x_3)}.$$

The three roots of Eq. (14) are explicitly given by

$$x_1 = \frac{1}{3\beta}\left(\beta^2 - \rho_e\beta + \rho_e^2\right),$$
$$x_2 = \frac{\beta}{6}\left(\rho_e^2 - \frac{2\rho_e}{\beta} - 1 + \sqrt{3}(1-\rho_e^2)i\right), \quad [16]$$
$$x_3 = \frac{\beta}{6}\left(\rho_e^2 - \frac{2\rho_e}{\beta} - 1 - \sqrt{3}(1-\rho_e^2)i\right),$$

where $\beta = \left(-\rho_e^3 + 27\bar{\kappa}^3/2 + 3\sqrt{3}\sqrt{27\bar{\kappa}^6/4 - \rho_e^3\bar{\kappa}^3}\right)^{1/3}$ and $\bar{\kappa} = (\alpha_2\rho_c\mu)^{1/3}$.

***5.2.1. Dominant balance.*** To simplify the above expressions for the eigenvalues of Eq. (13), we use the method of dominant balance (20). We note that $x_1$ is the largest positive root, while $x_2$ and $x_3$ have negative real part. As such, the time course of $O_0(t)$ will be quickly dominated by the exponential growing term $e^{x_1 t}$, as the terms $e^{x_2 t}$ and $e^{x_3 t}$ will decay to zero relative to $e^{x_1 t}$. For this reason, we can simplify the formula for $O_0(t)$ by keeping only the $e^{x_1 t}$ term while making sure that the initial condition $O_0(0) = 0$ is met:

$$O_0(t) \simeq C_1\left(e^{x_1 t} - 1\right). \quad [17]$$

To simplify our expression even further, we note that the exact formula for $x_1$ is asymptotically equal to in the limit of slow conversion and dissociation:

$$x_1 \simeq \left(2k_c k_+ k_{\text{oligo},2} m_{\text{tot}}^{n_{\text{oligo},2}+n_{\text{conv}}+1}\right)^{1/3} \equiv \bar{\kappa}. \quad [18]$$

This result Eq. (18) can be obtained by means of a dominant balance argument on the characteristic equation

$$x^3 + \rho_e x^2 - \alpha_2\rho_c\mu = 0 \quad [19]$$

in the limit $\rho_c, \rho_e \ll 1$, i.e. when $\rho_c = \bar{\rho}_c\varepsilon$, $\rho_e = \bar{\rho}_e\varepsilon$ with $\varepsilon$ small and $\bar{\rho}_c, \bar{\rho}_e = \mathcal{O}(1)$. The basic idea of this method is identify two terms in Eq. (19) that can be matched in such a way that all remaining terms in the equation vanish as $\varepsilon \to 0$. It is easy to see that the relevant dominant balance for our problem is obtained by matching the first and third term in Eq. (19), which suggests $x = \mathcal{O}(\varepsilon^{1/3})$. In fact, writing $x = \varepsilon^{1/3}X$ with $X = \mathcal{O}(1)$, we find

$$X^3 + \bar{\rho}_e\varepsilon^{2/3}X^2 - \alpha_2\bar{\rho}_c\mu = 0$$

Since the second term vanishes as $\varepsilon \to 0$, we can solve

$$X^3 - \alpha_2\bar{\rho}_c\mu = 0 \quad \Rightarrow \quad X \simeq (\alpha_2\bar{\rho}_c\mu)^{1/3}.$$

The largest eigenvalue of interest is therefore approximatively equal to

$$x \simeq (\alpha_2\rho_c\mu)^{1/3} = \bar{\kappa},$$

which proves Eq. (18). Note that in the opposing limit of fast conversion, $\rho_c, \rho_e \gg 1$, the dominant balance argument recovers $\kappa = (\alpha_2\mu)^{1/2}$, which is the rate of aggregate proliferation in the absence of oligomer conversion (Eq. (33)) (16).

***5.2.2. Early-time solutions.*** Using Eq. (18), we can simplify the relevant expression for $C_1$, which becomes $C_1 \simeq \frac{\alpha_1}{3\bar{\kappa}}$. Hence, we arrive at the following simple expression for the concentration of oligomers at early times:

$$O_0(t) \simeq \frac{\alpha_1}{3\bar{\kappa}}\left(e^{\bar{\kappa}t} - 1\right). \quad [20]$$

The accuracy of Eq. (20) against numerical solution of Eq. (11) is shown in Supplementary Fig. 12, which demonstrates that Eq. (20) is able to capture the early-time exponential increase of oligomer concentrations. Using Eq. (12b) and Eq. (12c) in combination with the expression for $O_0(t)$, Eq. (20), we obtain the early-time solutions for $P_0(t)$ and $M_0(t)$ respectively through simple integration. In summary, during the early stages of the aggregation reaction, the concentration of oligomers, fibril number and fibril mass all increase exponentially with time according to

$$O_0(t) = \frac{\alpha_1}{3\bar{\kappa}}\left(e^{\bar{\kappa}t} - 1\right) \quad [21a]$$

$$P_0(t) = \frac{\alpha_1 \rho_c}{3\bar{\kappa}^2}\left(e^{\bar{\kappa}t} - 1\right) \quad [21b]$$

$$M_0(t) = \frac{\bar{\lambda}^3}{3\bar{\kappa}^3}\left(e^{\bar{\kappa}t} - 1\right) \quad [21c]$$

with

$$\bar{\lambda} = \left(2k_c k_+ k_{\text{oligo},1} m_{\text{tot}}^{n_{\text{oligo},1}+n_{\text{conv}}}\right)^{1/3} \quad [21d]$$

$$\bar{\kappa} = \left(2k_c k_+ k_{\text{oligo},2} m_{\text{tot}}^{n_{\text{oligo},2}+n_{\text{conv}}+1}\right)^{1/3}. \quad [21e]$$

**5.3. A note on the effective proliferation rate of oligomer dynamics.** Through the mathematical analysis of the linearized equations Eq. (12), we demonstrated that, during the early stages of aggregation, the concentration of oligomers increases exponentially with time and we have identified that this exponential dynamics of oligomer concentrations is controlled by the following combination of the individual rate parameters

$$\bar{\kappa} = \left(2k_c k_+ k_{\text{oligo},2} m_{\text{tot}}^{n_{\text{oligo},2}+n_{\text{conv}}+1}\right)^{1/3}.$$

It is interesting to note that the effective rate $\bar{\kappa}$ is the geometric mean of the individual rates for oligomer formation, oligomer conversion and fibril elongation. In fact, $\bar{\kappa}$ can be written as

$$\bar{\kappa} = \left(2k_c k_+ k_{\text{oligo},2} m_{\text{tot}}^{n_{\text{oligo},2}+n_{\text{conv}}+1}\right)^{1/3} = (\alpha_2 \rho_c \mu)^{1/3}$$

where $\alpha_2 = k_{\text{oligo},2} m_{\text{tot}}^{n_{\text{oligo},2}}$ is the rate (units $s^{-1}$) of oligomer formation, $\rho_c = k_c m_{\text{tot}}^{n_{\text{conv}}}$ is the rate (units $s^{-1}$) of oligomer conversion and $\mu = 2k_+ m_{\text{tot}}$ is the rate (units $s^{-1}$) of fibril elongation. This result can be interpreted in terms of the Hinshelwood autocatalytic cycle (Supplementary Fig. 13(a)), which describes a series of $N$ first order chemical reactions that are connected in a circular manner (22, 23). A key feature of the Hinshelwood autocatalytic cycle is that the concentration of each species in the cycle increases in proportion to the concentration of the previous species in the cycle. Another essential feature of the Hinshelwood reaction system is that the $N$ reactions are connected in a cyclic fashion, i.e. that the rate of formation of the first species in the cycle, $X_1$ depends on the concentration of the last species in the cycle, $X_N$; this characteristic implies that the system behaves as a positive feedback loop. Indeed, Hinshelwood (23) showed that, after an initial phase of adaptation, the concentrations of all species in the cycle will increase exponentially with time. This exponential growth is controlled by a single effective rate which is the same for all species along the cycle and equal to the geometric mean of the constituent rates:

$$(k_1 k_2 \cdots k_N)^{1/N}. \quad [22]$$

where $N$ is the number of steps in the Hinshelwood cycle and $k_i$, $i = 1, 2, \cdots, N$ are the individual rates along the cycle. Oligomer dynamics in the linearized limit can be understood as a Hinshelwood autocatalytic cycle with $N = 3$ species: fibril mass ($M$) is responsible for generating oligomers ($O$); these oligomers in turn convert to elongation competent species ($P$); elongation increases the amount of fibril surface (mass) available for generating further oligomers through secondary nucleation. This is a Hinshelwood autocatalytic cycle because the rates of each step (oligomer formation, oligomer conversion and fibril elongation) depend on the concentrations $M$, $O$ and $P$, respectively. Moreover, the species are connected in a circular fashion, leading to a positive feedback loop (Supplementary Fig. 13(b)). The emergence of this autocatalytic cycle in the early-time dynamics explains why the effective proliferation rate for aggregates and oligomers is the geometric mean of the rates of oligomer formation, oligomer conversion and fibril growth, $\kappa = (\alpha_2 \rho_c \mu)^{1/3}$. Moreover, if the reaction rate $k_i$ order of each step is $n_i$, $i = 1, 2, \cdots, N$, the concentration dependence of the overall rate constant for growth will be the arithmetic mean of the individual reaction orders

$$\frac{n_1 + n_2 + \cdots + n_N}{N}. \quad [23]$$

Applied to the case of oligomer dynamics, Eq. (23) implies that the monomer concentration dependence of the overall aggregate proliferation rate $\bar{\kappa} \propto m_{\text{tot}}^{\gamma}$, will be the (arithmetic) mean of the reaction orders of oligomer formation, oligomer conversion and fibril elongation:

$$\begin{aligned}\gamma &= \frac{n_{\text{oligo},2} + n_{\text{conv}} + 1}{3} \\ &= \frac{(\text{reaction order of oligomer formation}) + (\text{reaction order of oligomer conversion}) + (\text{reaction order of elongation})}{3}\end{aligned} \quad [24]$$

**5.4. Self-consistent solution for fibril mass.** An accurate solution for $M(t)$ can be constructed by combining Eq. (21c) with the late-time behavior $M(\infty) = m_{\text{tot}}$ using a single function:

$$\frac{M(t)}{m_{\text{tot}}} = \frac{\frac{M_0(t)}{m_{\text{tot}}}}{1 + \frac{M_0(t)}{m_{\text{tot}}}} \simeq \frac{\frac{\bar{\lambda}^3}{3\bar{\kappa}^3}\left(e^{\bar{\kappa}t} - 1\right)}{1 + \frac{\bar{\lambda}^3}{3\bar{\kappa}^3}\left(e^{\bar{\kappa}t} - 1\right)}. \quad [25]$$

The performance of Eq. (25) against numerical solution of Eq. (11) is shown in Supplementary Fig. 14. It is interesting to note that the form of this solution is equivalent to Eq. (5) with the parameters in Eq. (33) being replaced as follows: $\kappa \to \bar{\kappa}$ and $\lambda^2/(2\kappa^2) \to \bar{\lambda}^3/(3\bar{\kappa}^3)$. The interpretation of the term $\bar{\lambda}^3/(3\bar{\kappa}^3)$ is the critical fibril mass necessary to start the autocatalytic cycle of secondary nucleation (oligomer formation, oligomer conversion and fibril elongation).

**5.5. Self-consistent solution for oligomer concentration.** To obtain a self-consistent solution for the oligomer concentration $O(t)$, we rewrite the defining dynamic equation (see Eq. (11))

$$\frac{dO(t)}{dt} = k_{\text{oligo},1} m(t)^{n_{\text{oligo},1}} + k_{\text{oligo},2} m(t)^{n_{\text{oligo},2}} M(t) - \rho_e O(t) \quad [26]$$

as an integral equation by formal integration using $O(0) = 0$. When $\rho_e$ is independent of monomer concentration, this yields the formal solution:

$$O(t) = e^{-\int_0^t \rho_e(t')dt'} \int_0^t e^{\int_0^\tau \rho_e(\tau')d\tau'} k_{\text{oligo},1} m(\tau)^{n_{\text{oligo},1}} d\tau + e^{-\int_0^t \rho_e(t')dt'} \int_0^t e^{\int_0^\tau \rho_e(\tau')d\tau'} k_{\text{oligo},2} m(\tau)^{n_{\text{oligo},2}} M(\tau) d\tau. \quad [27]$$

Since $\rho_e$ does not depend on time, we have $\int_0^t \rho_e(t')dt' = \rho_e t$, such that after using Eq. (25) in Eq. (27) and evaluating integrals explicitly, we arrive the following expression for the total concentration of unconverted oligomers:

$$O(t) = O_{\text{prim}}(t) + O_{\text{sec}}(t) \quad [28a]$$

with

$$O_{\text{prim}}(t) = \frac{e^{-\rho_e t}\bar{\lambda}^2}{2\rho_e(\alpha - 1)k_+\rho_c}\left[ {}_2F_1\left(1, 1 - n_1 + \frac{\rho_e}{\bar{\kappa}}, \frac{\rho_e}{\bar{\kappa}} + 1, \frac{\alpha}{\alpha - 1}\right) - e^{\rho_e t}\left(1 - \alpha + \alpha e^{\bar{\kappa}t}\right)^{1-n_1} {}_2F_1\left(1, 1 - n_1 + \frac{\rho_e}{\bar{\kappa}}, \frac{\rho_e}{\bar{\kappa}} + 1, \frac{\alpha}{\alpha - 1}e^{\bar{\kappa}t}\right)\right], \quad [28b]$$

and

$$O_{\text{sec}}(t) = \frac{e^{-\rho_e t}\bar{\kappa}^2}{2\rho_e(\alpha - 1)k_+\rho_c}\left[ {}_2F_1\left(1, 1 - n_2 + \frac{\rho_e}{\bar{\kappa}}, \frac{\rho_e}{\bar{\kappa}} + 1, \frac{\alpha}{\alpha - 1}\right) - {}_2F_1\left(1, -n_2 + \frac{\rho_e}{\bar{\kappa}}, \frac{\rho_e}{\bar{\kappa}} + 1, \frac{\alpha}{\alpha - 1}\right) - e^{\rho_e t}\left(1 - \alpha + \alpha e^{\bar{\kappa}t}\right)^{1-n_2} {}_2F_1\left(1, 1 - n_2 + \frac{\rho_e}{\bar{\kappa}}, \frac{\rho_e}{\bar{\kappa}} + 1, \frac{\alpha}{\alpha - 1}e^{\bar{\kappa}t}\right) + e^{\rho_e t}\left(1 - \alpha + \alpha e^{\bar{\kappa}t}\right)^{-n_2} {}_2F_1\left(1, -n_2 + \frac{\rho_e}{\bar{\kappa}}, \frac{\rho_e}{\bar{\kappa}} + 1, \frac{\alpha}{\alpha - 1}e^{\bar{\kappa}t}\right)\right], \quad [28c]$$

where ${}_2F_1(a, b, c, z)$ denotes the hypergeometric function, $\alpha = \bar{\lambda}^3/(3\bar{\kappa}^3)$. Supplementary Fig. 14, illustrates the performance of the solution Eq. (28) in comparison with numerics. Discrepancies between the two result from the non-exact form for $m(t)$ used but are lower than typical experimental error (see Supplementary Fig. 19).

**5.6. Unconverted and converted oligomers.** Supplementary Fig. 14 highlights that the concentration of (unconverted) oligomers, $O(t)$, is considerably greater than that of fibrillar oligomers (or converted oligomers, see Supplementary Sec. 4.3). Consequently, unconverted oligomers are the dominant contributor to oligomer populations. This fact can be shown as follows. When primary and secondary nucleation generate (unconverted) oligomers that need to undergo a conversion step before being able to grow further, the master equation Eq. (1) becomes:

$$\frac{\partial f(t,j)}{\partial t} = 2k_+ m(t) f(t, j-1) - 2k_+ m(t) f(t,j) + [k_c m(t)^{n_{\text{conv}}} + k_d] O(t) \delta_{j,n_2}, \quad [29]$$

where the terms on the second line of Eq. (29) describe the source of converted (or growth-competent) oligomers

$$\frac{dO(t)}{dt} = k_{\text{oligo},1} m(t)^{n_{\text{oligo},1}} + k_{\text{oligo},2} m(t)^{n_{\text{oligo},2}} M(t) - [k_c m(t)^{n_{\text{conv}}} + k_d] O(t). \quad [30]$$

Transforming Eq. (29) in the continuum limit we find

$$\frac{\partial f(t,x)}{\partial t} = -2k_+ m(t)\frac{\partial f(t,x)}{\partial x} + [k_c m(t)^{n_\text{conv}} + k_d]O(t)\delta(x-n_2), \qquad [31]$$

where $f(t,x)$ is the concentration of filaments of size $x$. Following the same procedure as for Eq. (7), we arrive at the following expression for the concentration of converted oligomers

$$f(t,n_2) = \frac{[k_c m(t)^{n_\text{conv}} + k_d]}{2k_+ m(t)}O(t). \qquad [32]$$

Thus, if the time evolution of $O(t)$ and $m(t)$ is known, then the full length distribution of aggregates in the continuum limit can be accessed. Moreover, plotting Eq. (32) in Supplementary Fig. 14 together with the concentration of unconverted oligomers clearly highlights that unconverted oligomers represent the dominant oligomeric species present at any time during the aggregation reaction.

**5.7. Primary vs secondary oligomers.** Using our kinetic framework and the rate parameters extracted from the experimental measurements of A$\beta$42 oligomer concentrations we can estimate the fraction of primary oligomers over the total concentration of oligomers (Supplementary Figs. 15 and 16). We find that primary oligomers are outnumbered by secondary oligomers.

### 6. Fitting of experimental data and kinetic parameter estimation

Fitting (and misfitting) of experimental data to the integrated rate laws developed in the previous sections is used to test different mechanistic hypotheses, to estimate quantitatively the various kinetic parameters and hence obtain a detailed mechanistic understanding of A$\beta$42 oligomer dynamics.

**6.1. Fibril mass concentration.** The first step in the kinetic analysis of experimental data consists in fitting kinetic data for the aggregate mass concentration recorded with various initial concentrations of monomeric protein (Figs. 2a,i, 2b,i and 2c,i).

***6.1.1. Mechanism 1.*** Normalized kinetic traces of fibril mass formation are fitted globally to Eq. (5) using the online available *Amylofit* platform (19). A global fit means that kinetic traces at various concentrations of monomers are fitted simultaneously using a single choice of the free parameters with the reaction orders $n_1$ and $n_2$ capturing any dependency on the monomer concentration (19). Note that in this case there are only two free parameters in the fitting of aggregate mass concentrations, which are $(k_+k_1)^{1/2}$, $(k_+k_2)^{1/2}$. The resulting fit is shown in Fig. 2a,i of the main text and the estimated kinetic parameters are: $(k_+k_1)^{1/2} = 30$ M$^{-1}$s$^{-1}$, $(k_+k_2)^{1/2} = 2.0 \times 10^5$ M$^{-\frac{3}{2}}$s$^{-1}$, $n_1 = 2$ and $n_2 = 2$, as reported previously in Ref. (16).

***6.1.2. Mechanisms 2 and 3.*** For mechanisms 2 and 3 normalized kinetic traces of fibril mass formation are fitted globally to Eq. (25) with the two free fitting parameters being $(k_+k_ck_\text{oligo,1})^{1/3}$ and $(k_+k_ck_\text{oligo,2})^{1/3}$. The resulting fits are shown in Figs. 2b,i and 2c,i of the main text and the estimated kinetic parameters are: $(k_\text{oligo,1}k_+k_c)^{1/3} = 7.3\times10^2$ M$^{-1}$s$^{-1}$, $(k_\text{oligo,2}k_+k_c)^{1/3} = 2.2\times10^5$ M$^{-\frac{3}{2}}$s$^{-1}$, $n_\text{oligo,1} + n_\text{conv} = 3.5$, $n_\text{oligo,2} + n_\text{conv} = 3.6$ (see Supplementary Table 1).

**6.2. Oligomer concentrations.**

***6.2.1. Mechanism 1.*** For Mechanism 1 the concentration of oligomers can be predicted solely on the basis of the fitting parameters for the aggregate mass using Eq. (9) and without introducing additional fitting parameters (see Supplementary Sec. 4.3). The concentration of oligomers over time predicted using this equation and the parameter values determined in Supplementary Sec. 6.1 is shown in Fig. 2a,ii of the main text.

***6.2.2. Mechanism 2.*** Oligomer concentration data are fitted to Eq. (28) with $k_d = 0$. The parameters $(k_+k_ck_\text{oligo,1})^{1/3}$ and $(k_+k_ck_\text{oligo,2})^{1/3}$, as well as the reaction orders $n_\text{oligo,1} + n_\text{conv}$ and $n_\text{oligo,2} + n_\text{conv}$, are known from the analysis of fibril mass concentration (Supplementary Sec. 6.1, Fig. 2b,i of the main text). Thus, using $k_+ = 3 \times 10^6$ M$^{-1}$s$^{-1}$ (16), there is only one free parameter that can be varied in the fit, namely $\rho_c$. Since the parameter $\rho_c$ controls both the oligomer peak height and the timescale for oligomer depletion, the best (mis)fit of our oligomer data to this model cannot explain the observed data (Fig. 2b,ii of the main text).

***6.2.3. Mechanism 3.*** Oligomer concentration data are fitted to Eq. (28). The parameters $(k_+k_ck_\text{oligo,1})^{1/3}$ and $(k_+k_ck_\text{oligo,2})^{1/3}$, as well as the reaction orders $n_\text{oligo,1} + n_\text{conv}$ and $n_\text{oligo,2} + n_\text{conv}$, are fixed by the analysis of fibril mass concentration (Supplementary Sec. 6.1). Thus, using $k_+ = 3 \times 10^6$ M$^{-1}$s$^{-1}$ (16), in the fitting of oligomer concentrations there are only two free parameters which can be varied. These are: $\rho_c$ and $\rho_e$. The best fit of our oligomer data at $m_\text{tot} = 5\mu$M is shown in Fig. 2c,i of the main text and Supplementary Fig. 19, and the extracted fitting parameters are summarized in Table 1. In Fig. 3a of the main text, we have used these fitting parameters to predict the time course of oligomer concentrations upon variation of the initial concentration of monomers, with the reaction orders $n_\text{oligo,1}$, $n_\text{oligo,2}$ and $n_\text{conv}$ capturing any dependency on the monomer concentration.

**6.3. Summary of extracted rate parameters.** We now summarize the values of the rate parameters extracted from the combined fits of fibril mass and oligomer concentrations using model 3:

$$\begin{aligned} k_+ &= 3 \times 10^6 \ \mathrm{M}^{-1}\mathrm{s}^{-1} \\ k_{\mathrm{oligo},1} &= 6.7 \times 10^{-8} \ \mathrm{M}^{0.2}\mathrm{s}^{-1} \\ k_{\mathrm{oligo},2} &= 2.0 \ \mathrm{M}^{-0.9}\mathrm{s}^{-1} \\ k_c &= 1.9 \times 10^9 \ \mathrm{M}^{-2.7}\mathrm{s}^{-1} \text{ (assumed to be the same for primary and secondary oligomers)} \\ k_e &= 9.7 \times 10^{-5} \ \mathrm{s}^{-1} \\ n_{\mathrm{conv}} &= 2.7 \text{ (assumed to be the same for primary and secondary oligomers)} \\ n_{\mathrm{oligo},2} &= 0.9 \\ n_{\mathrm{oligo},1} &= 0.8. \end{aligned}$$

From these parameters we can determine the reaction rates (units 1/s) for the different processes at $m_{\mathrm{tot}} = 5\ \mu\mathrm{M}$ (summarized in Fig. 4 of main text): $\alpha_1/m_{\mathrm{tot}} = k_{\mathrm{oligo},1} m_{\mathrm{tot}}^{n_{\mathrm{oligo},1}-1} = 7.7 \times 10^{-7}\ \mathrm{s}^{-1}$, $\alpha_2 = k_{\mathrm{oligo},2} m_{\mathrm{tot}}^{n_{\mathrm{oligo},2}} = 3.3 \times 10^{-5}\ \mathrm{s}^{-1}$, $\rho_c = k_c m_{\mathrm{tot}}^{n_{\mathrm{conv}}} = 9.4 \times 10^{-6}\ \mathrm{s}^{-1}$, $\rho_d = 8.8 \times 10^{-5}\ \mathrm{s}^{-1}$, $\mu = 2k_+ m_{\mathrm{tot}} = 30\ \mathrm{s}^{-1}$ (Supplementary Fig. 18 and Supplementary Table 2).

To obtain the rates for the primary oligomers, we have assumed for simplicity that their conversion rate is the same as that of the secondary oligomers. This may not be the case, however, and is not directly testable with the current data: although both primary and secondary pathways contribute to the ensemble of oligomers, the large majority of species are formed by the latter (see Supplementary Figs. 15 and 16).

**6.4. Kinetic analysis in the presence of Brichos.** Fibril mass data in the presence (Fig. 2d of the main text) or in the presence (Fig. 2e of the main text) of Brichos were fitted to Eq. (25) with $\bar{\kappa}$ and $\bar{\lambda}^3/(2\bar{\kappa}^3)$ as fitting parameters. This procedure yields: $\bar{\kappa}(\text{no Brichos}) = (2.3 \pm 0.2) \times 10^{-3}\ \mathrm{s}^{-1}$, and $\bar{\lambda}^3/(2\bar{\kappa}^3)(\text{no Brichos}) = (2.8 \pm 2.1) \times 10^{-4}$ for the data in Fig. 2d of the main text; $\bar{\kappa}(\text{Brichos}) = (4.7 \pm 0.2) \times 10^{-4}\ \mathrm{s}^{-1}$ and $\bar{\lambda}^3/(2\bar{\kappa}^3)(\text{no Brichos}) = (1.9 \pm 0.3) \times 10^{-2}$ for the data in Fig. 2e of the main text. The time courses of oligomer concentrations were then predicted using Eq. (28) and that differences in and affect only the rates of oligomer formation but keep the rates of conversion and dissociation fixed.

**6.5. Connection with previous studies of A$\beta$42 aggregation.** Using kinetic analysis we have determined the values for the reaction rates and reaction orders of key molecular steps of A$\beta$42 oligomer dynamics, including oligomer formation and oligomer conversion. How do these results relate to previous studies of A$\beta$42 fibril formation (see e.g. Ref (16))? In either case, a key result from the master equation analysis is the emergence of a dominant combined kinetic parameter that sets the fundamental timescale over which the overall aggregation process occurs. The specific form of this dominant kinetic parameter depends on the specific mechanism of fibril replication under consideration. In particular, previous studies of A$\beta$42 fibril formation have described secondary nucleation as a single-step process coupled to fibril elongation (mechanism 1). Kinetic analysis in this limit shows that the dominant kinetic parameter is

$$\kappa = \left(2k_+ k_2 m_{\mathrm{tot}}^{n_2+1}\right)^{1/2}, \qquad [33]$$

where $k_2 m_{\mathrm{tot}}^{n_2}$ is the rate of secondary nucleation (viewed as a single-step process) and $2k_+ m_{\mathrm{tot}}$. Note that this result makes perfect sense in view of the analogy with the Hinshelwood cycle (Supplementary Fig. 13); indeed, fibril self-replication occurs in two stages (secondary nucleation and elongation) and the dominant kinetic parameter $\kappa$ is nothing but the geometric average of the rates of these two steps.

In the present study, we have been able to decompose the overall secondary nucleation process into its elementary constituents: oligomer formation, oligomer conversion and fibril elongation (mechanism 3). In this case, we have found that the characteristic timescale of aggregation is

$$\bar{\kappa} = \left(2k_c k_+ k_{\mathrm{oligo},2} m_{\mathrm{tot}}^{n_{\mathrm{oligo},2}+n_{\mathrm{conv}}+1}\right)^{1/3}. \qquad [34]$$

In this case, a complete fibril self-replication step occurs in three stages (oligomer formation, oligomer conversion and fibril elongation) and indeed the dominant kinetic parameter $\bar{\kappa}$ is simply the geometric average of the rates of these three processes. It is important to note that, since the aggregation of fibrils is characterized by a single (measurable) timescale, the parameters $\kappa$ and $\bar{\kappa}$ must be equal. In other words, the characteristic timescale for amyloid fibril aggregation (and its dependence on monomer concentration, $\kappa, \bar{\kappa} \propto m_{\mathrm{tot}}^{\gamma}$) are independent of whether we describe fibril self-replication as a two- or three-step process. In previous studies of A$\beta$42 amyloid fibril formation (16), secondary nucleation is described as a single-step process that generates fibrillar oligomers. Thus, the autocatalytic cycle associated with secondary nucleation consists of two steps (secondary nucleation and elongation) and the overall (measurable) rate of fibril proliferation is the geometric average of the rates associated with these two steps (Supplementary Fig. 17):

$$\kappa = (k_2 m_{\mathrm{tot}}^{n_2} \cdot 2k_+ m_{\mathrm{tot}})^{1/2} = \left(3 \times 10^{-7}\ \mathrm{s}^{-1} \cdot 30\ \mathrm{s}^{-1}\right)^{1/2} \simeq 0.003\ \mathrm{s}^{-1}.$$

In this study (mechanism 3), direct measurements of oligomer concentrations over time have allowed us to ‘zoom’ into the cycle of secondary nucleation with ‘higher resolution’ and conclude that this cycle consists at least of 3 steps (oligomer formation, conversion, and elongation). The overall (measurable) rate of fibril proliferation is the geometric average of these three rates and yields the same numerical result as in (16) (Supplementary Fig. 17):

$$\bar{\kappa} = \left(k_{\text{oligo},2} m_{\text{tot}}^{n_{\text{oligo},2}} \cdot k_c m_{\text{tot}}^{n_{\text{conv}}} \cdot 2k_+ m_{\text{tot}}\right)^{1/3} = \left(3 \times 10^{-5}\ \text{s}^{-1} \cdot 9 \times 10^{-6}\ \text{s}^{-1} \cdot 30\ \text{s}^{-1}\right)^{1/3} \simeq 0.002\ \text{s}^{-1}.$$

Moreover, using the reaction orders for (secondary) oligomer formation, oligomer conversion and fibril elongation determined in this study, we obtain that the overall concentration dependence of $\bar{\kappa} \propto m_{\text{tot}}^{\gamma}$ is

$$\gamma = \frac{n_{\text{oligo},2} + n_{\text{conv}} + 1}{3} = \frac{0.9 + 2.7 + 1}{3} = 1.5. \tag{35}$$

This value is the same as the one determined by combining the reaction orders of secondary nucleation (viewed as a single-step process) and elongation determined in Ref. (16), such that $\kappa \propto m_{\text{tot}}^{\gamma}$ with:

$$\gamma = \frac{n_2 + 1}{3} = \frac{2 + 1}{2} = 1.5.$$

Thus, the cycle of oligomer formation, oligomer conversion and fibril elongation determined in this study generates an effective rate of fibril proliferation $\bar{\kappa}$ and a scaling exponent $\gamma$ that is consistent with previous studies (16).

The equality between $\kappa$ and $\bar{\kappa}$ implies a direct relationship between the reaction rates and reaction orders of oligomer formation, oligomer conversion and fibril elongation determined in this study and the reaction rates and reaction orders for secondary nucleation (single-step) and elongation determined previously in Ref. (16). The rate constant for single-step secondary nucleation is

$$k_2 = \left(\frac{k_{\text{oligo},2}^2 k_c^2}{2k_+}\right)^{1/3} \tag{36}$$

with reaction order

$$n_2 = \frac{2n_{\text{oligo},2} + 2n_{\text{conv}} - 1}{3}. \tag{37}$$

Moreover, the expression for the aggregate mass concentration in the presence of an oligomer conversion step is given by (see Eq. (25))

$$\frac{M(t)}{m_{\text{tot}}} = \frac{\frac{\bar{\lambda}^3}{3\bar{\kappa}^3}\left(e^{\bar{\kappa}t} - 1\right)}{1 + \frac{\bar{\lambda}^3}{3\bar{\kappa}^3}\left(e^{\bar{\kappa}t} - 1\right)}. \tag{38}$$

When secondary nucleation is described as a single step (Ref. (16)), the corresponding expression for the aggregate mass reads

$$\frac{M(t)}{m_{\text{tot}}} = \frac{\frac{\lambda^2}{2\kappa^2}\left(e^{\kappa t} - 1\right)}{1 + \frac{\lambda^2}{2\kappa^2}\left(e^{\kappa t} - 1\right)}. \tag{39}$$

where

$$\lambda = (2k_+ k_1 m_{\text{tot}}^{n_1})^{1/2}. \tag{40}$$

Since Eq. (38) and Eq. (39) describe the same measurable quantify, we must have

$$\frac{\bar{\lambda}^3}{3\bar{\kappa}^3} = \frac{\lambda^2}{2\kappa^2}. \tag{41}$$

This condition yields a direct relationship between the reaction rates and reaction orders of oligomer formation by primary nucleation, oligomer conversion and fibril elongation determined in this study and the reaction rates and reaction orders for primary nucleation (single-step) and elongation determined previously in Ref. (16) assuming a single-step nucleation process. The rate constant for single-step primary nucleation is

$$k_1 = \frac{2}{3}\left(\frac{k_{\text{oligo},1}^3 k_c^2}{2k_+ k_{\text{oligo},2}}\right)^{1/3}. \tag{42}$$

Moreover, assuming that $n_{\text{conv}}$ is the same for primary and secondary oligomers, we find for the reaction orders

$$n_1 = n_{\text{oligo},1} - n_{\text{oligo},2} + n_2. \tag{43}$$

We can verify that the fitting parameters determined in this study (Supplementary Table 1) together with the relationships Eq. (36), Eq. (37), Eq. (42) and Eq. (43) are consistent with the rate parameters for single-step primary and secondary nucleation determined previously in Ref. (16)

$$k_2 = \left(\frac{k_{\text{oligo},2}^2 k_c^2}{2k_+}\right)^{1/3} = \left(\frac{(2\ \text{M}^{-0.9}\ \text{s}^{-1})^2(1.9 \times 10^9\ \text{M}^{-2.7}\ \text{s}^{-1})^2}{2 \times 3 \times 10^6\ \text{M}^{-1}\ \text{s}^{-1}}\right)^{1/3} \simeq 1.3 \times 10^4\ \text{M}^{-2}\ \text{s}^{-1}$$

$$k_1 = \frac{2}{3}\left(\frac{k_{\text{oligo},1}^3 k_c^2}{2k_+ k_{\text{oligo},2}}\right)^{1/3} = \frac{2}{3}\left(\frac{(6.7 \times 10^{-8}\ \text{M}^{0.2}\ \text{s}^{-1})^3(1.9 \times 10^9\ \text{M}^{-2.7}\ \text{s}^{-1})^2}{2 \times (3 \times 10^6\ \text{M}^{-1}\ \text{s}^{-1}) \times (2\ \text{M}^{-0.9}\ \text{s}^{-1})}\right)^{1/3} \simeq 3.0 \times 10^{-4}\ \text{M}^{-1}\ \text{s}^{-1}$$

and

$$n_2 = \frac{2n_{\text{oligo},2} + 2n_{\text{conv}} - 1}{3} = \frac{2 \times 0.9 + 2 \times 2.7 - 1}{3} \simeq 2.1$$
$$n_1 = n_{\text{oligo},1} - n_{\text{oligo},2} + n_2 \simeq 2.0.$$

### 7. Supplementary Figures

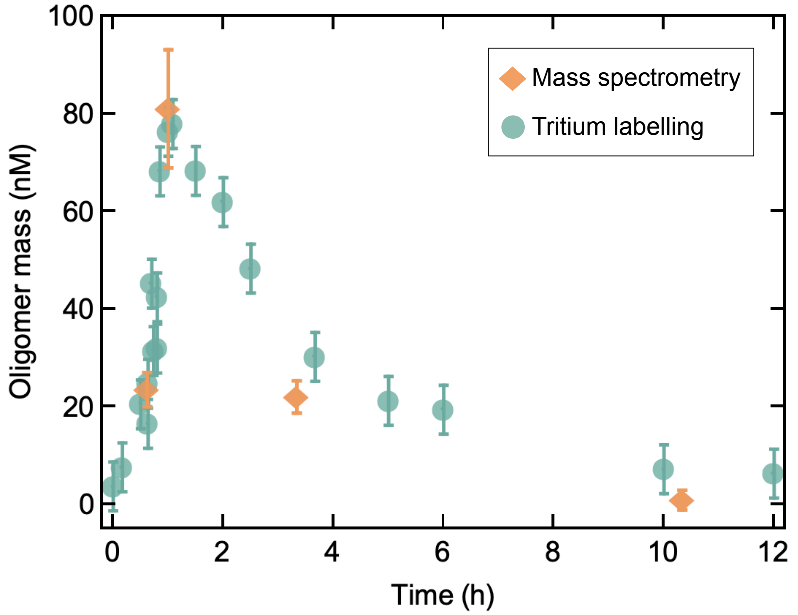

**Fig. 1.** Experimental time course of A$\beta$42 oligomer concentrations for $m_{\rm tot} = 5\mu$M in 20 mM sodium phosphate buffer, 0.2 mM EDTA, pH 8.0, 37$^o$C, recorded using mass spectrometry (see Supplementary Sec. 1.7) (orange diamonds) and comparison with the oligomer concentrations measurements performed using tritium labelling for the same monomer concentration (see Supplementary Sec. 1.6) (green circles). The two independent measurements of A$\beta$42 oligomer concentrations agree within experimental error.

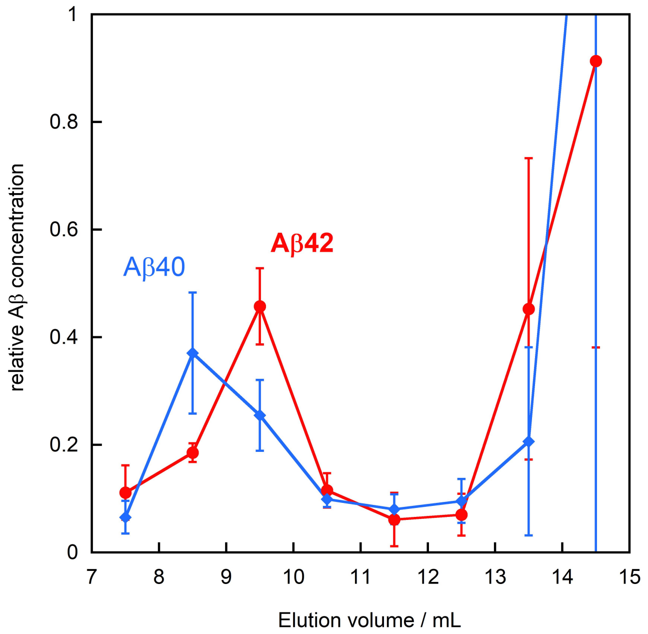

**Fig. 2.** Fractions of A$\beta$42 and A$\beta$40 oligomers as a function of elution volume from SEC (Supplementary Sec. 1.7). We note that A$\beta$40 oligomers are somewhat larger than A$\beta$42 oligomers, as they elute earlier from the SEC column. Experimental conditions as described in Supplementary Sec. 1.7 and in the caption to Fig. 1 of the main text. The relative A$\beta$ concentration in each fraction was then calculated as concentration divided by the summed concentration over fractions 7-12, and we show here the average and standard deviation over all time-points examined. Therefore the monomer peak (monomer elutes in fraction 14-16, only 14 and 15 analysed to show where it starts) can have a value above 1.

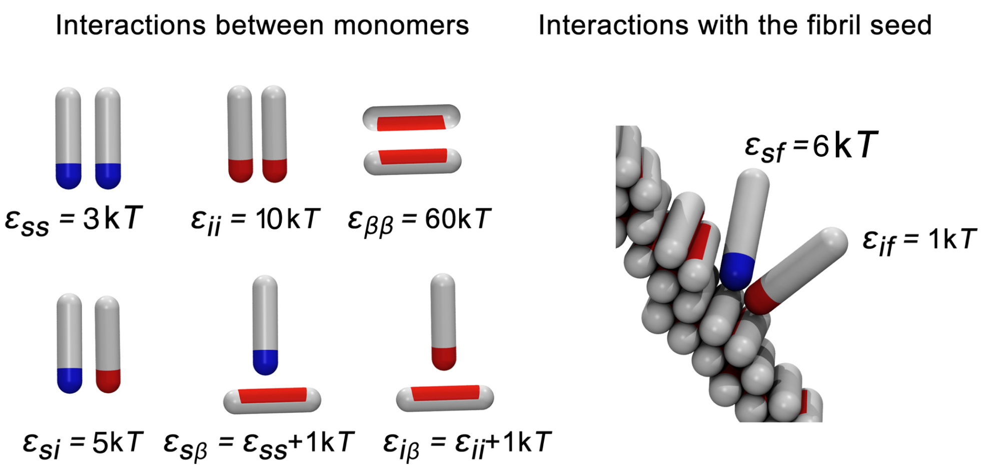

**Fig. 3.** Computer model: possible interactions in the system, and their values (in units of thermal energy $kT$). The protein exists in three states: monomeric ('s'), oligomer-forming ('i') and fibril-forming ('$\beta$'). This leads to 6 interaction parameters. $\varepsilon_{\beta\beta}$ is the strongest of these interactions, leading to effectively irreversible fibril formation. Monomers in the s and i states interact with the fibril surface; this controls monomer adsorption at the fibril surface and the subsequent detachment of sufficiently large oligomers.

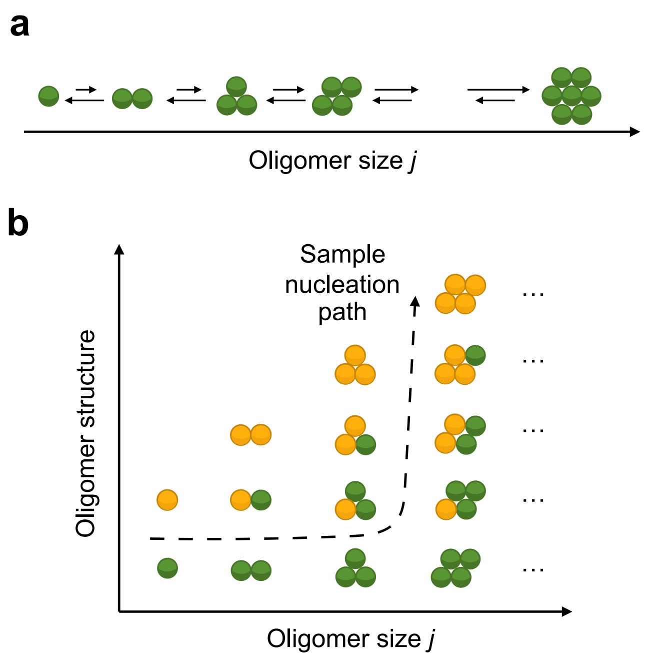

**Fig. 4.** (a) Classical nucleation theory involves aggregate size as the only relevant degree of freedom. (b) Two-step (or multi-step) nucleation involves additional degrees of freedom associated with aggregate structure (in our case $\beta$-sheet content). Adapted from (9).

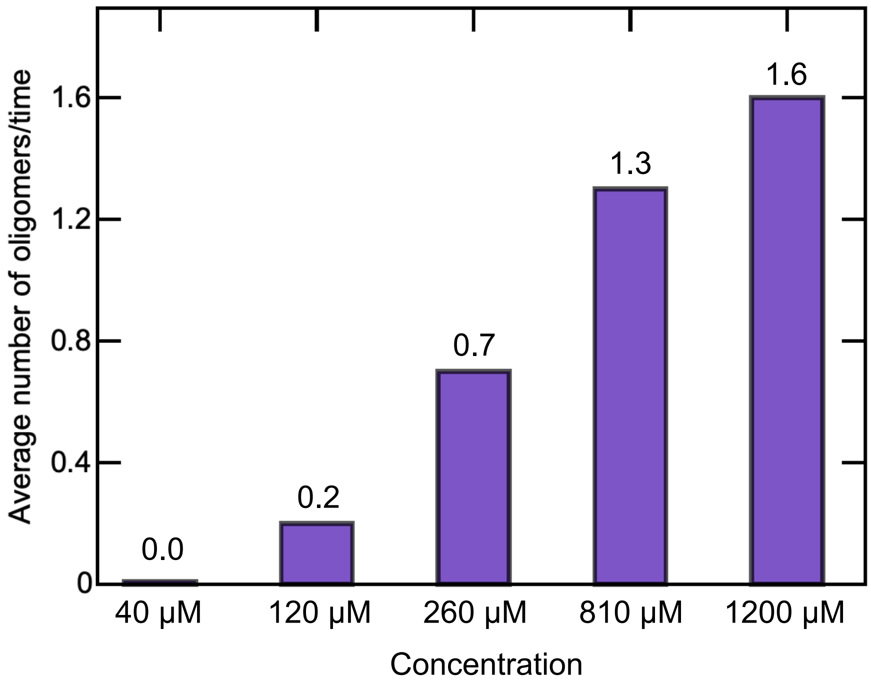

**Fig. 5.** Average number of oligomers in the system increases with the increase in monomer concentration.

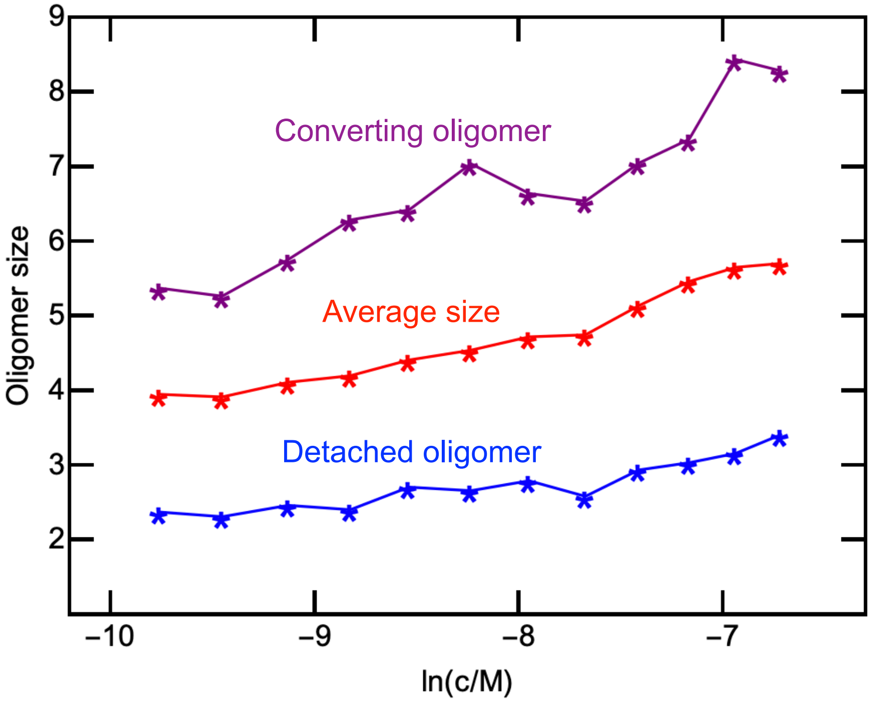

**Fig. 6.** The average size of oligomers after they detach from the fibril surface, the average oligomer size, and the size of converting oligomers as a function of monomer concentration. Note that the reaction order for conversion measured in the simulations ($n_{\mathrm{conv}} = 2$) does not correspond to the actual size of converting oligomers.

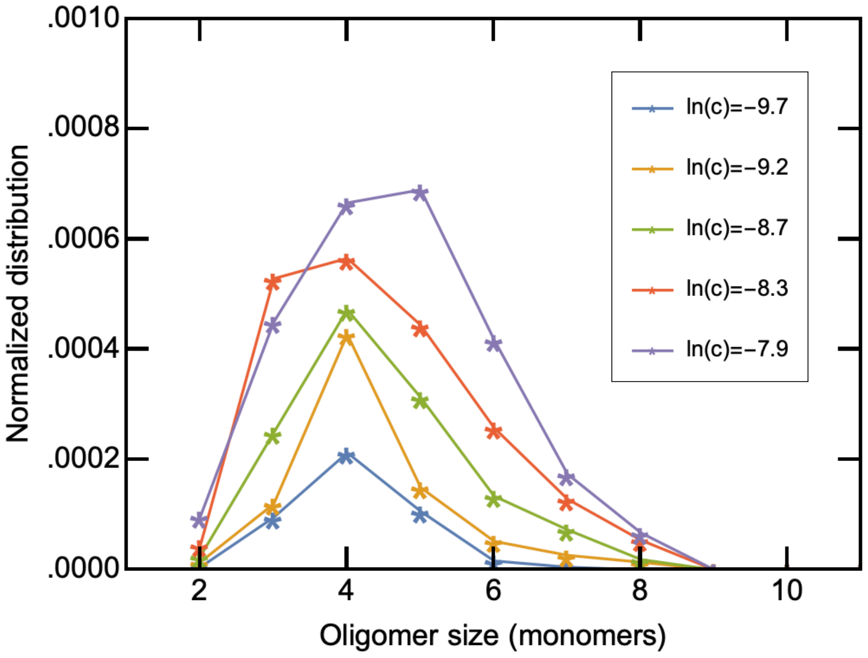

**Fig. 7.** Size distribution of oligomers obtained from computer simulations for different monomer concentrations.

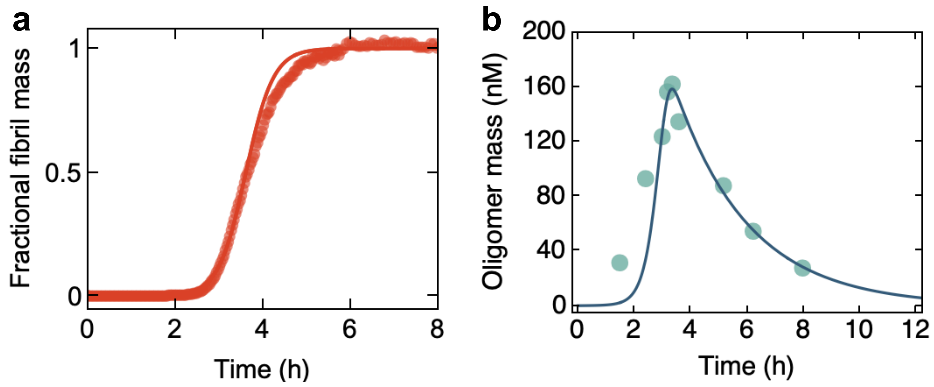

**Fig. 8.** Experimental time course of (a) fibril mass fraction and (b) oligomer concentrations for the length variant A$\beta$40 at $m_{\rm tot} = 10\mu$M in 20 mM sodium phosphate buffer, 0.2 mM EDTA, pH 7.4, 37$^o$C. The solid lines indicate fits to $\mathrm{Eq.}$ $(25)$ and $\mathrm{Eq.}$ $(28)$. The fits yield the following values for the rates of oligomer conversion and dissociation: $\rho_c \simeq 1 \times 10^{-6}$ s$^{-1}$ and $\rho_e \simeq 1 \times 10^{-4}$ s$^{-1}$. Experimental conditions as described in the caption to Fig. 1 of the main text.

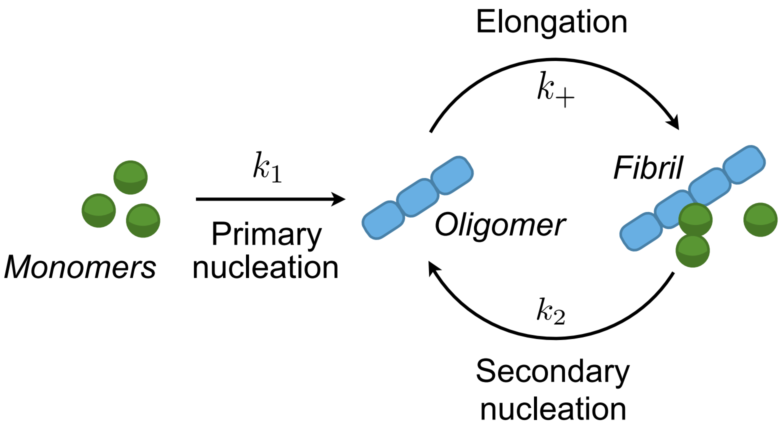

**Fig. 9.** Schematic representation of mechanism 1: oligomers generated by primary and secondary nucleation are short fibrillar aggregates which are able to grow into mature fibrils by recruiting free monomers.

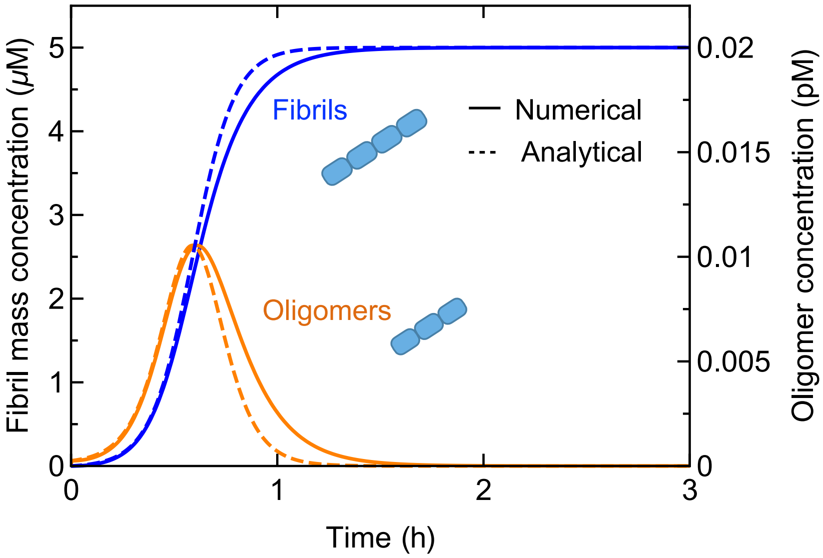

**Fig. 10.** Time course of fibril mass and oligomer concentrations predicted for mechanism 1 (solid lines: numerical solution to Eq. (1); dashed lines: analytical solutions, Eq. (5) and Eq. (9)). The calculation parameters are the rate constants extracted from the fitting of the aggregate mass data of Fig. 2a,i of the main text: $m_{\mathrm{tot}} = 5\ \mu$M, $k_+ = 3 \times 10^6$ M$^{-1}$s$^{-1}$, $k_1 = 3 \times 10^{-4}$ M$^{-1}$s$^{-1}$, $k_2 = 1.3 \times 10^5$ M$^{-2}$s$^{-1}$, $n_1 = n_2 = 2$ (16).

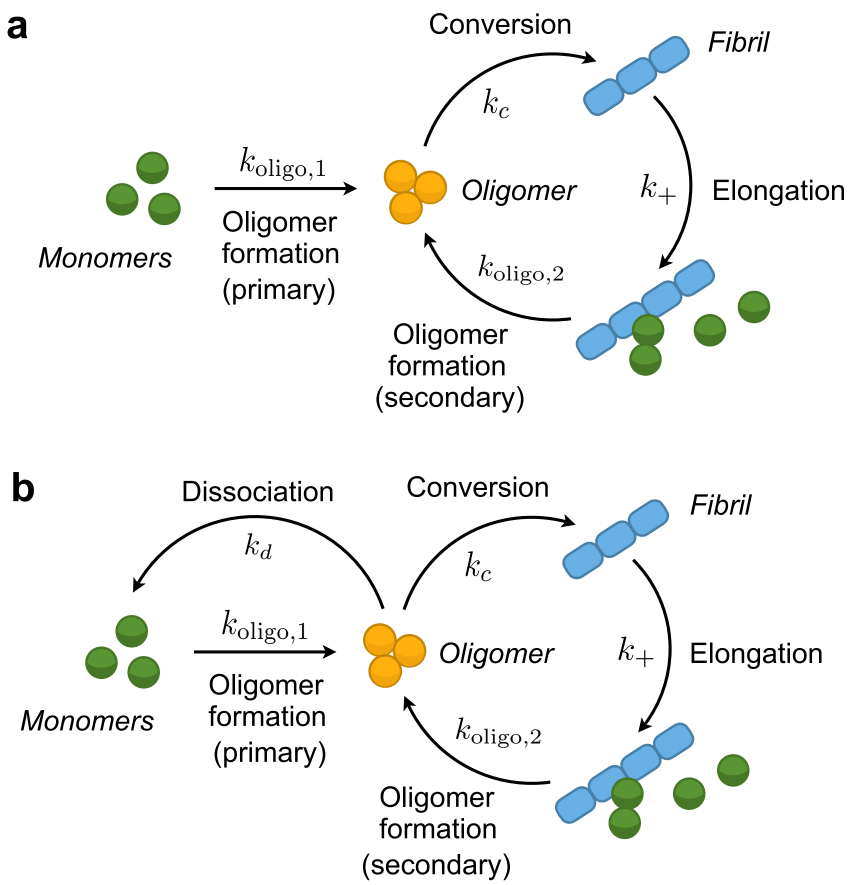

**Fig. 11.** Schematic representation of (a) mechanism 2 and (b) mechanism 3. These correspond to the mechanisms in Figs. 2b and 2c of the main text.

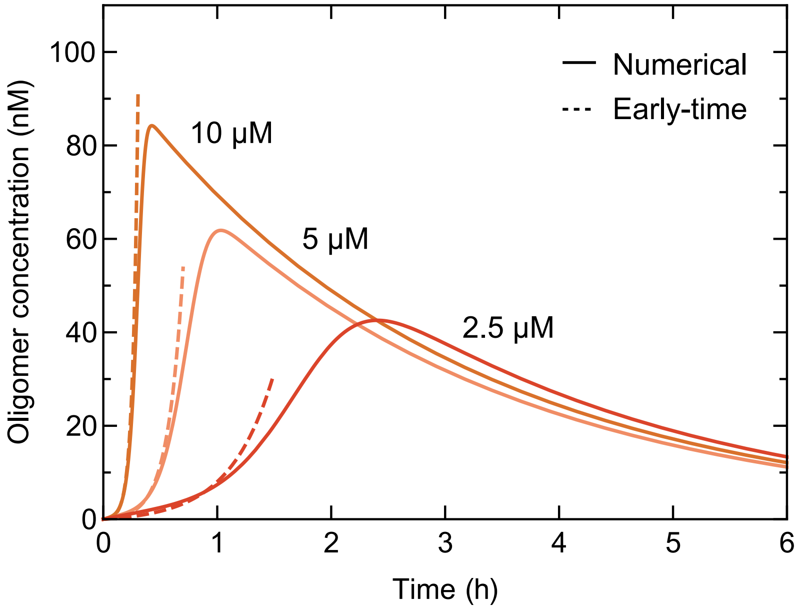

**Fig. 12.** Comparison between numerical solution of the kinetic equations, Eq. (11), (solid lines) and the early-time solution for oligomer concentration, Eq. (20) (dashed lines) for three distinct monomer concentrations. The curves were generated predicted using the fitting parameters of Supplementary Table 1.

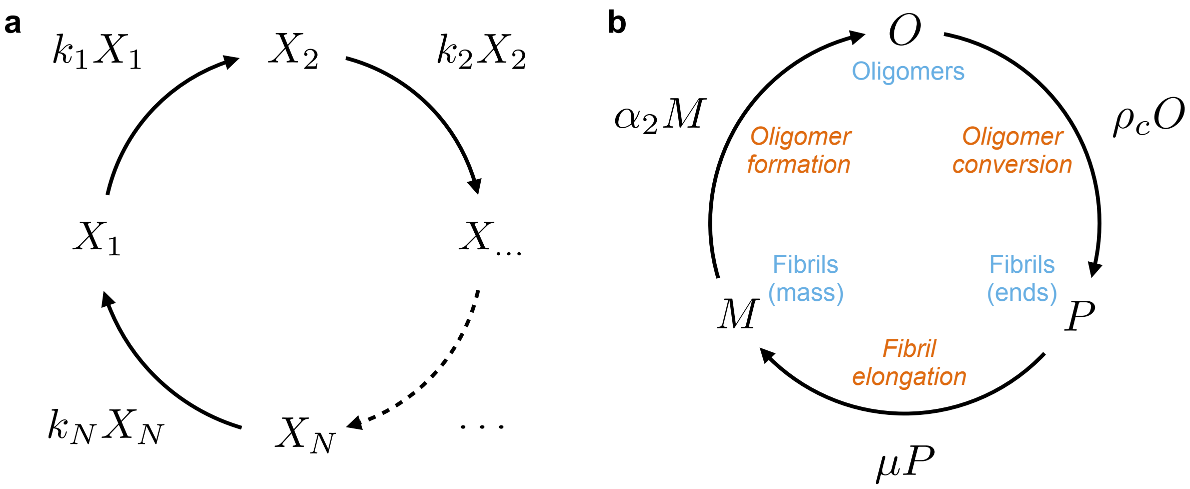

**Fig. 13.** (a) Schematic representation of a general Hinshelwood autocatalytic cycle (23). After sufficient time, the concentration of all species in the cycle increases exponentially with time with an effective rate of proliferation that is the geometric mean of the individual rates along the cycle, $(k_1k_2\cdots k_N)^{1/N}$. (b) Autocatalytic cycle for secondary nucleation. Oligomer dynamics in the linearized limit can be understood as a Hinshelwood autocatalytic cycle with $N=3$ species: fibril mass $(M)$, oligomer concentration $(O)$ and fibril number concentration $(P)$. The direct consequence of this observation is that the effective proliferation rate for aggregates and oligomers is the geometric mean of the rates of oligomer formation, oligomer conversion and fibril growth, $\kappa=(\alpha_2\rho_c\mu)^{1/3}$.

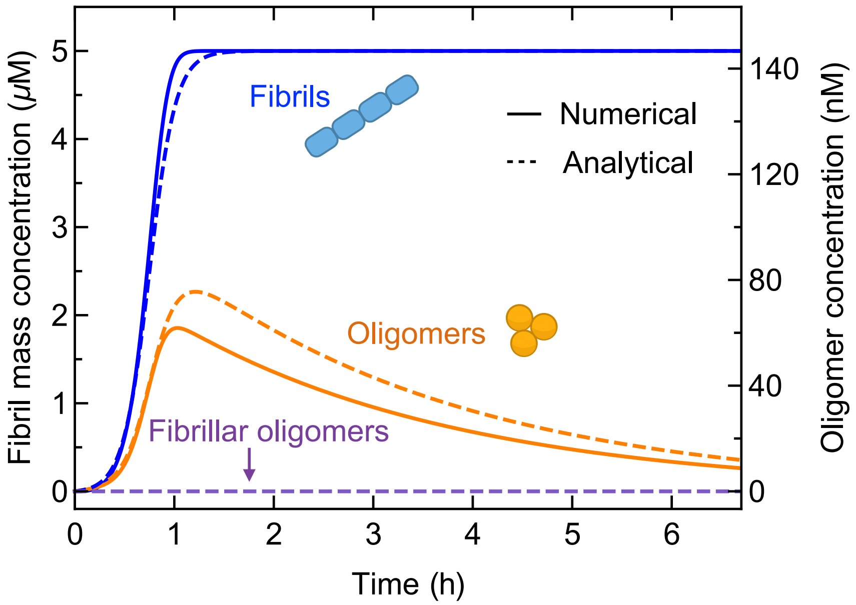

**Fig. 14.** Time evolution of fibril mass, $M(t)$, and oligomers, $O(t)$, predicted by the analytical solutions, $\mathrm{Eq.}\ (25)$ and $\mathrm{Eq.}\ (28)$, (dashed lines) is compared with the numerical solution of $\mathrm{Eq.}\ (11)$ (solid lines). The curves generated using the fitting parameters of Supplementary Table 1 and Supplementary Sec. 6. The figure illustrates also the predicted concentration of fibrillar oligomers (converted oligomers, $\mathrm{Eq.}\ (32)$) as a function of time.

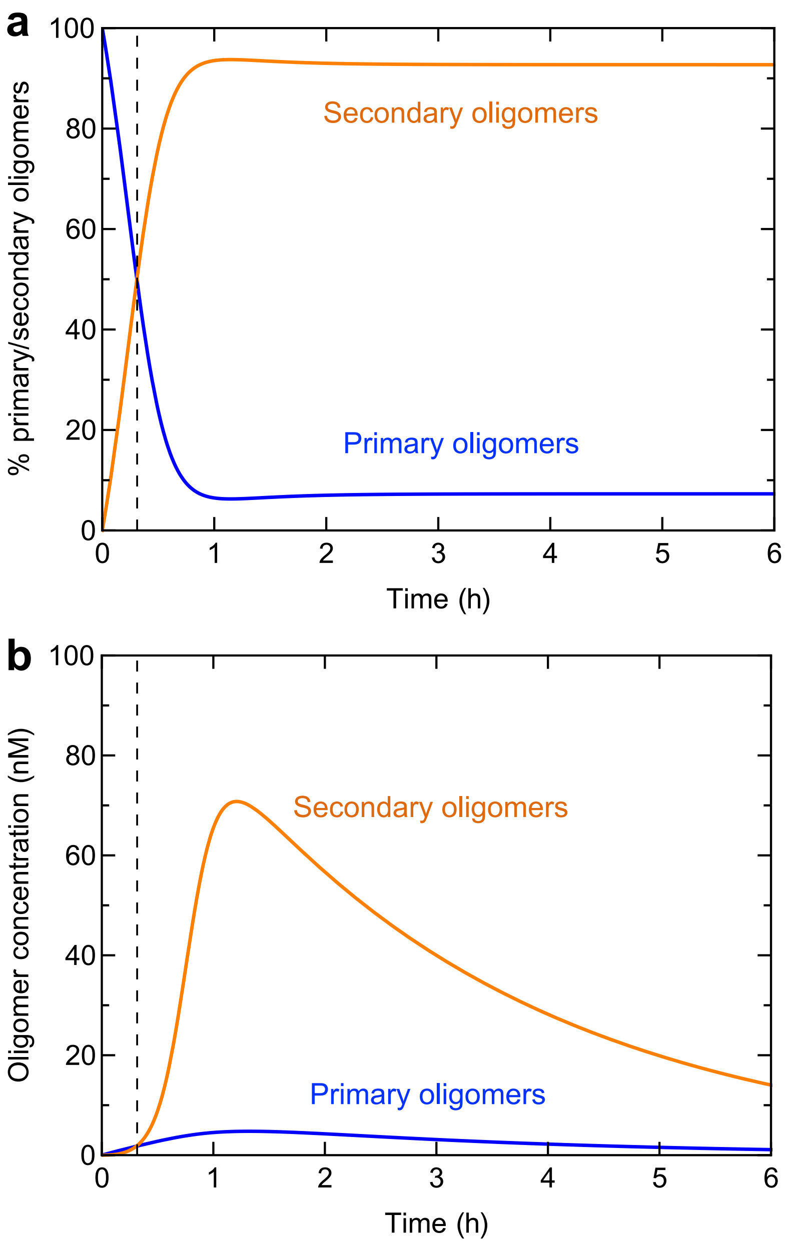

**Fig. 15.** Time evolution of the (a) fractions and (b) concentrations of primary (orange) and secondary (blue) oligomers during an ongoing aggregation reaction with an initial monomer concentration of 5 $\mu$M (predicted using the fitting parameters of Supplementary Table 1 and Supplementary Sec. 6). The figure also displays the predicted concentration of converted oligomers (short fibrils) as a function of time.

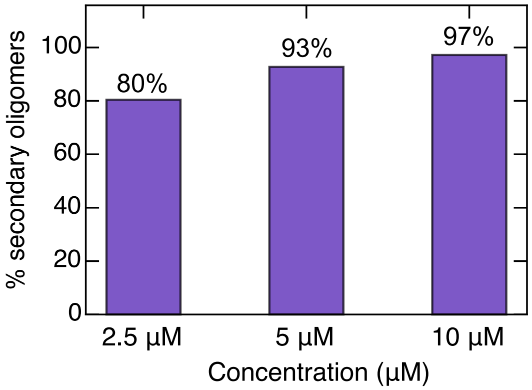

**Fig. 16.** Steady-state fractions of secondary oligomers as a function of monomer concentration (predicted using the fitting parameters of Supplementary Table 1 and Supplementary Sec. 6).

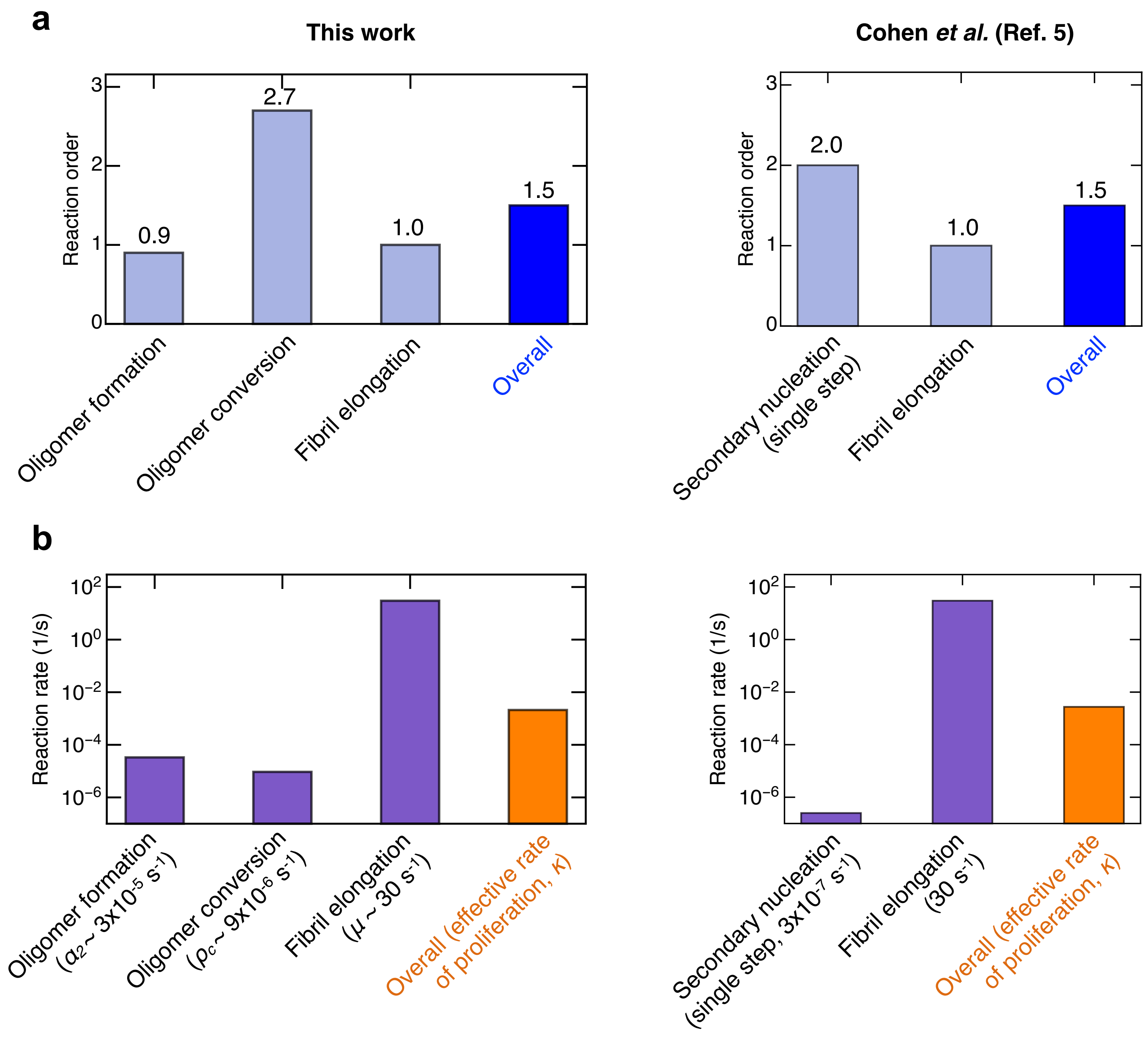

**Fig. 17.** Relationship between the reaction orders (a) and reaction rates (b) determined in this study and previous studies of A$\beta$42 aggregation (Ref. (16)). See Supplementary Sec. 6.5 for a discussion.

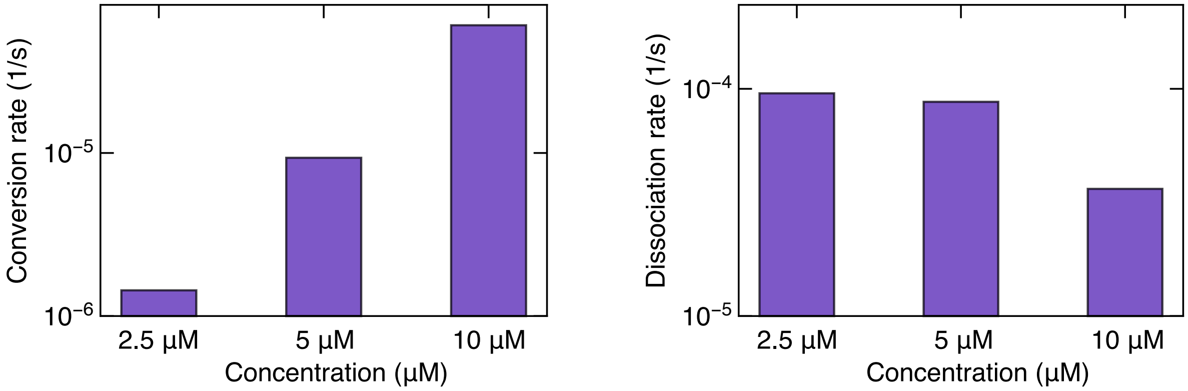

**Fig. 18.** Conversion and dissociation rates of oligomers as a function of monomer concentration (Supplementary Table 2).

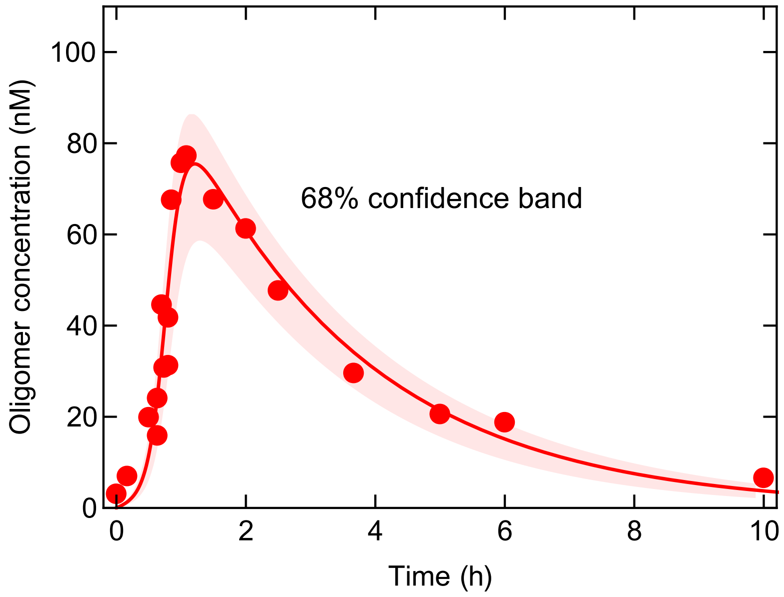

**Fig. 19.** Experimental time course of oligomer concentration recorded in this study for $m_{\rm tot} = 5\mu$M is fitted using model 3 (Eq. (28)). The shaded area represents the 68% confidence band.

### 8. Supplementary Tables

**Table 1. Summary of fitting parameters (using $k_+ = 3 \times 10^6$ M$^{-1}$s$^{-1}$)**

| Parameter | Estimate | Standard error | Obtained from fitting of |
|---|---|---|---|
| $(k_{\text{oligo},1} k_+ k_c)^{1/3}$ | $7.3 \times 10^2$ M$^{-1}$s$^{-1}$ | $1.1 \times 10^2$ M$^{-1}$s$^{-1}$ | Fibril mass concentration |
| $(k_{\text{oligo},2} k_+ k_c)^{1/3}$ | $2.2 \times 10^5$ M$^{-\frac{3}{2}}$s$^{-1}$ | $2 \times 10^4$ M$^{-\frac{3}{2}}$s$^{-1}$ | Fibril mass concentration |
| $\rho_e$ | $9.7 \times 10^{-5}$ s$^{-1}$ | $1.2 \times 10^{-5}$ s$^{-1}$ | Oligomer concentration |
| $\rho_c$ | $9.3 \times 10^{-6}$ s$^{-1}$ | $5 \times 10^{-7}$ s$^{-1}$ | Oligomer concentration |

**Table 2. Oligomer conversion and dissociation rates as a function of monomer concentration (using $k_+ = 3 \times 10^6$ M$^{-1}$s$^{-1}$)**

| Monomer concentration | 2.5 $\mu$M | 5 $\mu$M | 10 $\mu$M |
|---|---|---|---|
| Conversion rate $\rho_c$ | $1.4 \times 10^{-6}$ s$^{-1}$ | $9.3 \times 10^{-6}$ s$^{-1}$ | $6.1 \times 10^{-5}$ s$^{-1}$ |
| Dissociation rate $\rho_d$ | $9.7 \times 10^{-5}$ s$^{-1}$ | $8.8 \times 10^{-5}$ s$^{-1}$ | $3.6 \times 10^{-5}$ s$^{-1}$ |
| Fraction of converting oligomers $\frac{\rho_c}{\rho_c+\rho_d}$ | 1.4% | 9.7% | 63% |

### 9. Supplementary Movie

Excerpt from a simulation of secondary nucleation showing monomers (colored in blue) adsorbing onto the fibril (colored in white) and detaching as oligomers (colored in red), which dissolve several times before converting into a new fibril nucleus capable of growing (in white).
